# Altered microglial communication in brains with Alzheimer’s Disease pathology

**DOI:** 10.64898/2026.08.13.744723

**Authors:** Loren dos Santos, Ricardo A. Vialle, Natacha Comandante-Lou, Gilad Green, Yanling Wang, Shinya Tasaki, Naomi Habib, Vilas Menon, Philip L. De Jager, Roberto T. Raittz, David A. Bennett, Katia de Paiva Lopes

## Abstract

Microglia are the resident immune cells of the central nervous system and are highly versatile, continuously monitoring the brain microenvironment with their motile processes and responding to perturbations in diverse ways. Recent studies have revealed substantial functional heterogeneity among microglial states, suggesting that analyzing individual subpopulations is essential to capture specialized signaling programs and disease-associated interactions that would otherwise be obscured in bulk analyses. To better understand how these cells interact, we performed a ligand-receptor communication analysis using single-nucleus RNA sequencing data from the dorsolateral prefrontal cortex, comprising 16 microglial subpopulations together with other myeloid cells identified in brain tissue, including macrophages and monocytes. We characterized the signaling roles of these cell populations, compared individuals with and without neuropathologically defined Alzheimer’s disease, and associated cell-pair interactions with AD-related clinical and neuropathological traits. We found that AD was characterized by altered microglial composition and rewiring of intercellular communication, including disease-specific signaling pathways and distinct interaction hubs. Lipid-associated microglia were associated with tau pathology, whereas nuclear receptor signaling microglia were linked to hippocampal sclerosis. These findings were replicated in three independent datasets. Overall, this work provides an initial but comprehensive map of microglial communication in the aging human brain and identifies candidate signaling interactions for future functional validation.

## INTRODUCTION

Microglia are the resident macrophages of the central nervous system (CNS) and serve as its primary immune cells (Ajami et al. 2007; Arcuri et al. 2017). Although they account for a small proportion of brain cells (Mittelbronn et al. 2001), they play essential roles in maintaining brain homeostasis, including immune surveillance, phagocytosis, synaptic remodeling, neurodevelopment, and neurogenesis (Tan et al. 2020; Gao et al. 2023). Rather than existing in a single resting state, microglia exhibit remarkable functional plasticity, adopting distinct transcriptional and metabolic programs depending on brain region, age, sex, and local environmental cues (Young et al. 2021; Paolicelli et al. 2022; Lopes et al. 2022). These diverse functional states are coordinated through cell-cell communication, in which ligand-receptor (L-R) interactions and downstream signaling pathways regulate how microglia sense and respond to changes in their microenvironment (Armingol et al. 2021).

Microglia have emerged as important players in Alzheimer’s disease (AD), the leading cause of dementia. Genetic studies have shown that many AD risk variants regulate genes predominantly expressed in microglia, including *TREM2*, *CD33*, and *APOE* (Raj et al. 2014; Lambert et al. 2013; Bellenguez et al. 2022). During disease progression, the accumulation of amyloid-β plaques, hyperphosphorylated tau tangles, and other damage-associated molecular patterns (DAMPs) triggers reactive microglial states that can initially limit pathology but may ultimately promote chronic neuroinflammation and neurodegeneration (Heneka et al. 2015; De Strooper and Karran 2016). Advances in single-cell transcriptomics have identified multiple disease-associated microglial states with distinct transcriptional signatures (Keren-Shaul et al. 2017; Grubman et al. 2019; Mathys et al. 2019; Green et al. 2024). However, it remains unexplored whether these disease-associated shifts in microglial composition (e.g. decrease of homeostatic and increase of damage-associated states) result from changes in cell abundance or are also influenced by extensive reorganization of intercellular L-R communication.

In this study, we characterized microglial cell-cell communication in the dorsolateral prefrontal cortex (DLPFC) using single-nucleus RNA sequencing (snRNA-seq) data from 437 ROSMAP participants. We reconstructed L-R interaction networks across 16 microglial subpopulations as well as brain-associated macrophages and infiltrating monocytes, identified signaling pathways associated with AD pathology, and investigated how communication between specific microglial states relates to neuropathological and clinical phenotypes. To evaluate the robustness of our findings, we replicated these analyses in three independent snRNA-seq datasets (480 additional participants), yielding a total of 917 participants across four studies and validating our findings across diverse genetic backgrounds (**Figure 1**).

**Figure 1:**
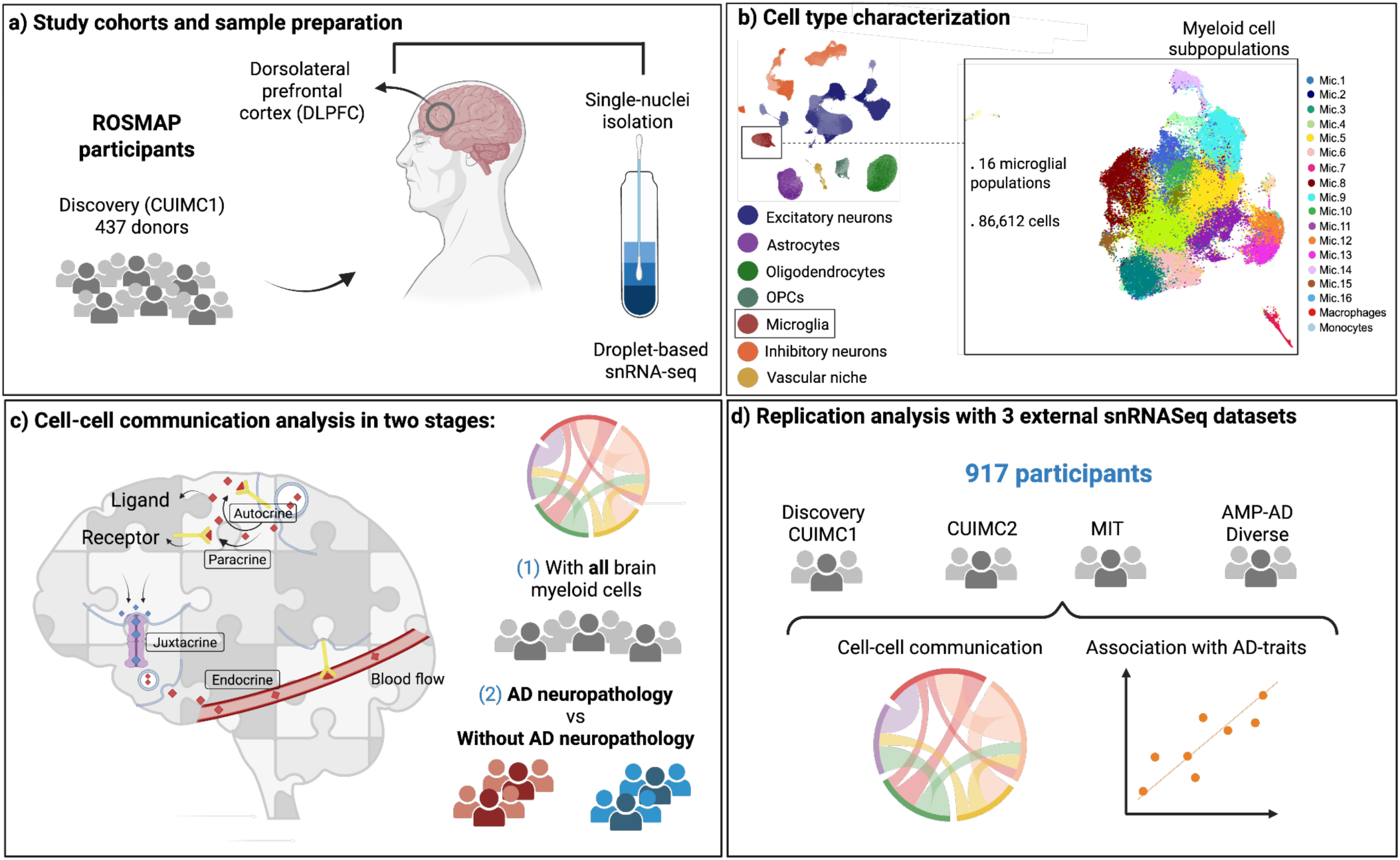
Study overview. **a)** Participants in the ROSMAP study cohorts and sample preparation. **b)** Clustering and annotation of microglia cell groups and subpopulations - Data obtained from Green et al. Nature, 2024. **c)** Analysis of cell communication (1) including cells from all participants to generate a comprehensive map of L-R interactions across the entire DLPFC microglial repertoire and (2) performing analyses separately between AD and non-AD cases to highlight differences in cell communication in the context of the disease. **d)** Replication of Discovery results with association analyses between pairs of myeloid cells and AD-related phenotypes.

## RESULTS

### Characteristics of the participants

The snRNA-seq data from the 437 individuals comprising the Discovery (Columbia University Irving Medical Center - CUIMC1) set are derived from participants in two longitudinal clinical pathological cohorts of aging and dementia: the Religious Orders Study (ROS) and the Rush Memory and Aging Project (MAP), commonly referred to as ROSMAP. Both studies were approved by the Institutional Review Board of Rush University Medical Center (Chicago, IL). The participants consisted of 219 older adults from MAP and 218 from ROS who were dementia free at enrollment and agreed to participate in annual clinical evaluations and organ donation. Participants were enrolled at a mean age of 81.2 years (SD: 7.1) and died at a mean age of 89.2 years (SD: 6.8), of whom 67.5% were female. At the time of death, 34.3% had no cognitive impairment, 25.4% had mild cognitive impairment (MCI), and the remaining 40.3% had dementia. Overall, 26.3% of participants carried at least one *APOE* ε4 allele. Upon autopsy, 62.7% received a pathological diagnosis of AD according to the NIA-Reagan criteria. TDP-43 pathology, extending beyond the amygdala, was observed in 30.9% of participants. Cerebrovascular diseases, including macroscopic infarcts and microinfarcts, were identified in 37.1% and 25.6% of cases, respectively. Additionally, moderate to severe amyloid angiopathy was observed in 34.9% of cases, and atherosclerosis and arteriolosclerosis were found in 40.0% and 38.7% of brains, respectively (**Table 1** and **Supplementary Table 1**).

**Table 1.** Description of the participants in the Discovery dataset.

| Description | Characteristics of the study participants |
| --- | --- |
| Participants (n) | 437 |
| Study (n) | MAP (n=219), ROS (n=218) |
| Age baseline, mean (SD), y | 81.2 (7.1) |
| Age at death, mean (SD), y | 89.2 (6.8) |
| Female sex, No. (%) | 295 (67.5) |
| Educational level, mean (SD), y | 16.3 (3.5) |
| Length of follow-up, mean (SD), y | 7.2 (4.6) |
| MMSE score, median (IQR) - at baseline | 28 (27.0-29.0) |
| Global cognition score, mean (SD) - at baseline | -0.09 (0.63) |
| Global cognition score, mean (SD) - at death | -0.86 (1.18) |
| MCI, No. (%) | 111 (25.4) |
| Dementia, No. (%) | 176 (40.3) |
| NIA-Reagan AD, No. (%)<sup>a</sup> | 274 (62.7) |
| Global AD pathologic score, median (IQR) | 0.64 (0.17-1.13) |
| Neuritic plaques score, median (IQR) | 0.64 (0.02-1.30) |
| Diffuse plaques score, median (IQR) | 0.54 (0.05-1.23) |
| Neurofibrillary tangles score, median (IQR) | 0.37 (0.15-0.71) |
| Macroscopic infarcts, No. (%) | 162 (37.1) |
| Microinfarcts, No. (%) | 112 (25.6) |
| Neocortical Lewy bodies, No. (%) | 35 (8.3) |
| TDP-43, No. (%)<sup>b</sup> | 128 (30.9) |
| Hippocampal sclerosis, No. (%) | 32 (7.4) |
| Amyloid angiopathy, No. (%)<sup>c</sup> | 150 (34.9) |
| Atherosclerosis, No. (%)<sup>c</sup> | 174 (40.0) |
| Arteriolosclerosis, No. (%)<sup>c</sup> | 168 (38.7) |
| APOE ε4, No. (%) | 115 (26.3) |
| <sup>a</sup> Intermediate or high likelihood. |  |
| <sup>b</sup> Inclusion beyond the amygdala. |  |
| <sup>c</sup> Moderate or severe. |  |

### A cell-cell interaction map of the prefrontal cortex myeloid cells

To construct a comprehensive map of interactions between brain myeloid cells, we performed cell communication analysis using snRNA-seq data from the DLPFC. The dataset consists of 84,064 previously annotated cells from 437 ROSMAP participants, grouped into 18 myeloid populations (16 microglial subpopulations, macrophages, and monocytes; **Figure 2a** and **Supplementary Table S2**). Using the literature-curated Cellchat database, we retained 2,425 highly variable L-R interactions across the myeloid populations, from the 3,234 validated molecular interactions available in the database (**Supplementary Figure S1**). Because there are different combinations of L-R pairs, these interactions resulted in 17,523 significant inferred cell-cell connections (permutation p < 0.05). At the unique L-R pair level, these connections involved 242 L-R pairs distributed across 74 signaling pathways (**Supplementary Tables 3-5**). While communication occurred broadly across all myeloid populations (**Supplementary Figures S2–S4**), Inflammatory microglia (Mic.15) emerged as the dominant signaling source, with 1,615 outgoing interactions, whereas Stress-response microglia (Mic.11) acted as the primary signaling target, receiving 3,474 incoming interactions (**Figure 2b** and **Supplementary Figure S5**). Importantly, these patterns were not explained by differences in overall cell abundance or transcript detection, as Mic.15 and Mic.11 were among the least abundant microglial populations and did not show increased numbers of detected genes compared with other microglial states (**Supplementary Table S6**).

**Fig 2.**
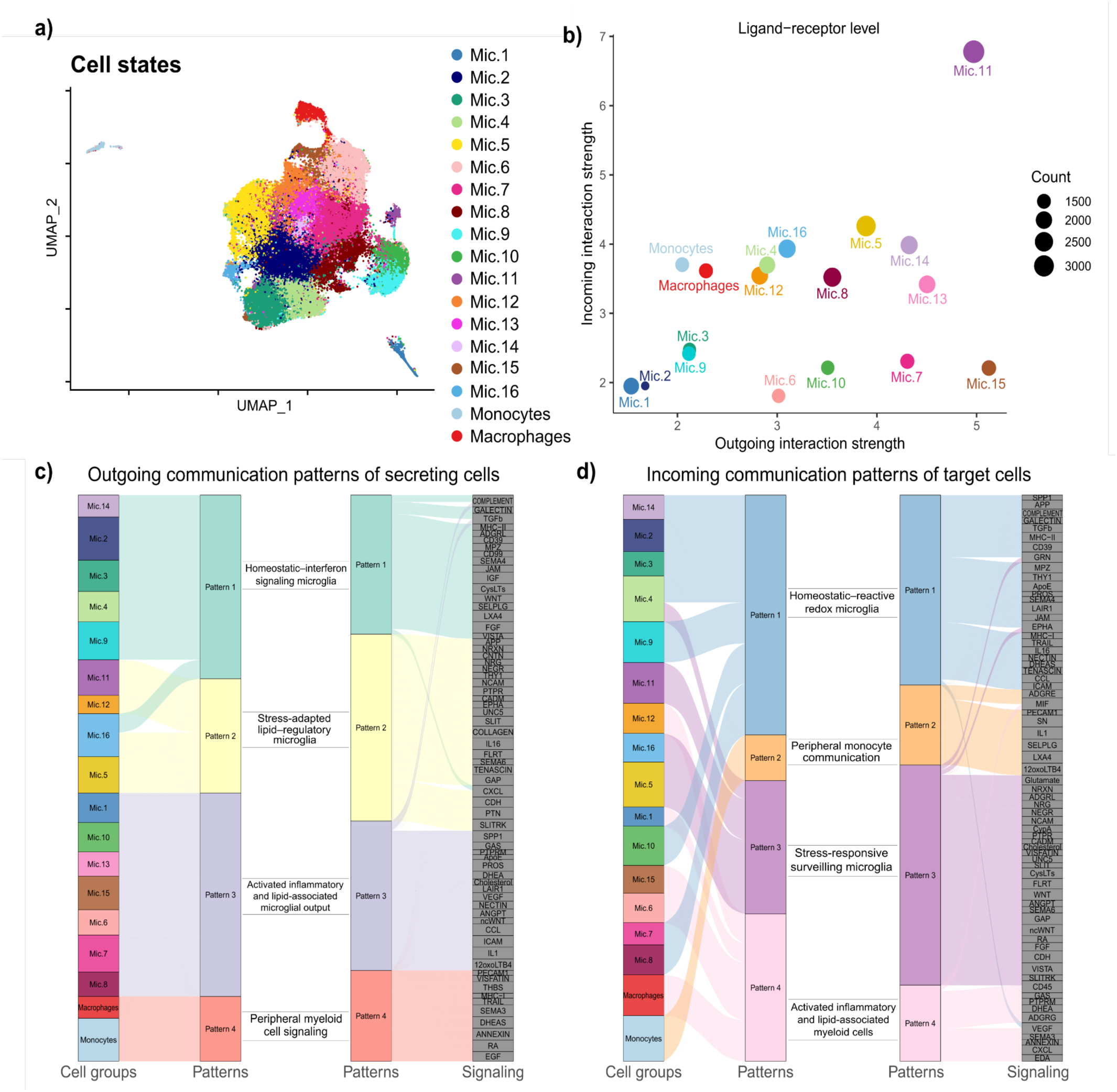
All brain myeloid cells. **a)** UMAP colored by cell states. **b)** Comparison of outgoing interaction strength (x-axis) and incoming interaction strength (y-axis) for cell communication results. The size of the dots is proportional to the number of L–R interactions. Colors are mapped according to panel (a). **c-d)** River plot showing outgoing and incoming cell groups and the most enriched signaling pathways associated with each inferred pattern, cutoff 0.4 for the NMF clusterization (see Methods).

To investigate higher-order communication programs, we applied non-negative matrix factorization (NMF) to identify coordinated outgoing and incoming signaling patterns (**Supplementary Tables 7-8**). Four outgoing and four incoming communication patterns were identified, distinguishing homeostatic, inflammatory, lipid-associated, stress-responsive, and peripheral myeloid cell signaling programs (**Figures 2c,d** and **Supplementary Figures S6-S9**). Analysis of pathway activity showed that outgoing signaling was relatively consistent across cell groups, whereas incoming signaling was more heterogeneous, with Mic.11 receiving signals through most pathways while Mic.15 displayed more pathway-specific incoming communication (**Supplementary Figure S10**). The ADGRE pathway exhibited uniformly strong outgoing activity across cell groups but strong incoming activity only in monocytes, whereas Glutamate, SPP1, APP, NRXN, TGFβ, and ADGRL showed cell-type-specific differences, particularly in Mic.11 and Mic.15. Finally, functional clustering grouped the 74 signaling pathways into five modules with distinct communication probability profiles, highlighting coordinated signaling programs across the myeloid interaction network (**Supplementary Figure S11**).

These findings reveal a complex and coordinated myeloid communication network, characterized by an apparent onset of inflammatory and stress responses in the aging human cortex.

### Altered signaling across microglial states in AD

To compare cell-cell communication across brains with and without AD pathology, we performed cell communication analyses separately on individuals with intermediate or high likelihood of AD according to the NIA–AA criteria (n = 274) and controls (CT) with low likelihood or no AD (n = 163) (**Supplementary Figure S12a** and **Supplementary Tables 9-10**). The AD group exhibited 16,204 interactions and a total interaction strength of 56.31 versus 13,212 interactions and a strength of 50.52 in CT (**Supplementary Figure S12b**). These differences were explained by the total number of cells in each group rather than AD status (two-tailed permutation tests P = 0.16 and P = 0.33, respectively, **Supplementary Figure S12d,e**). In addition, the observed fractions of subpopulations varied between groups. For instance, the Stress-response cells of Mic.11 and the Lipid-associated cells of Mic.13 were enriched in AD, while the Interferon response cells in Mic.14 and the Reacting cells of Mic.8 were observed in higher fractions in CT compared to the expected overall (**Figure 3a,b**). Such conditions are important when interpreting the cell communication results since some of the interactions can be entangled with the cell imbalance.

**Fig 3.**
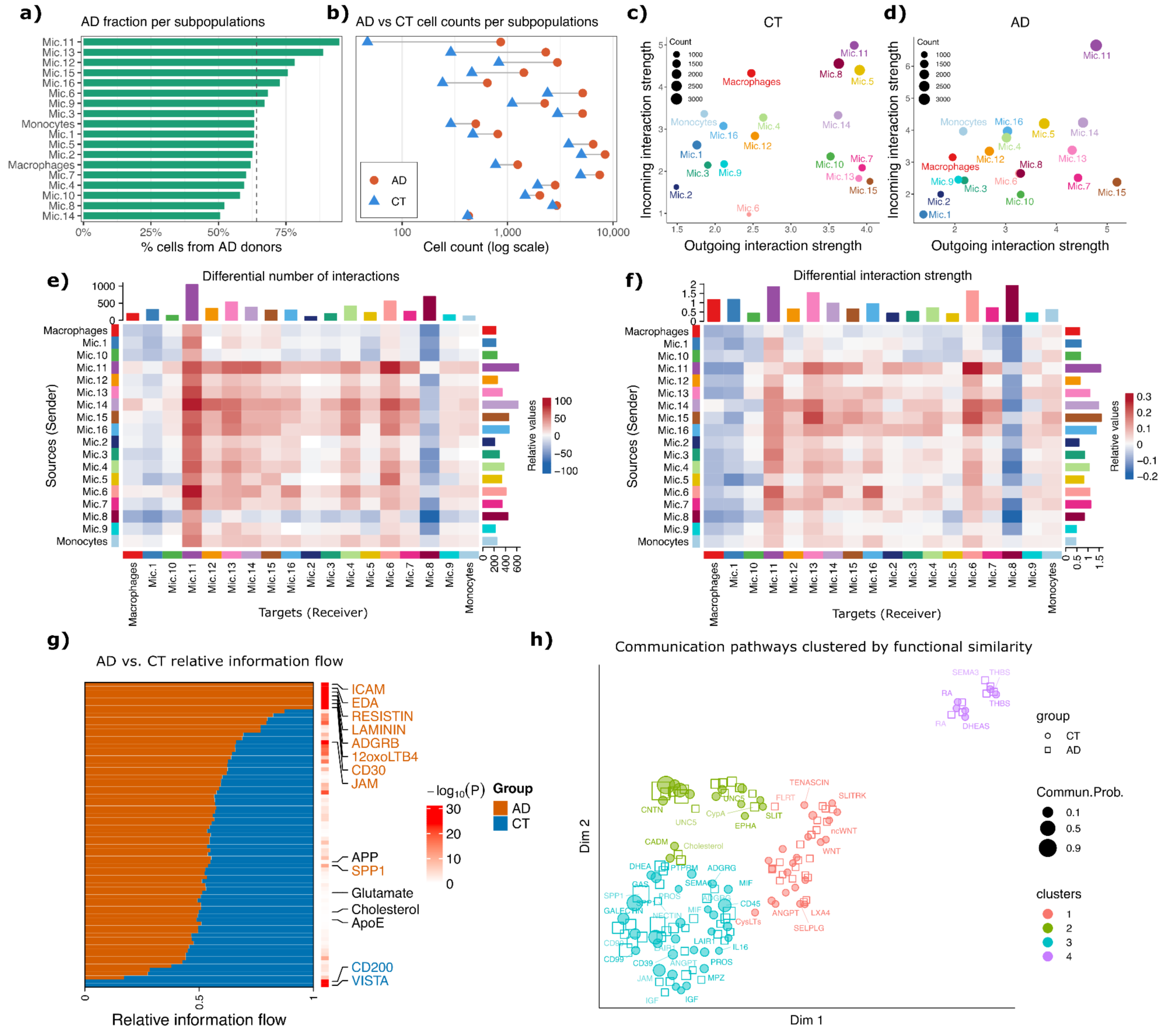
Differences in microglial cell-cell communication between AD and CT. **a)** Proportion of cells derived from AD participants across myeloid subpopulations, showing variable AD representation among cell states. **b)** Cell counts per myeloid subpopulation in AD and CT participants, demonstrating generally higher cell counts in AD across subpopulations. **c)** Scatter plot of outgoing interaction strength (x-axis) and incoming interaction strength (y-axis) for CT participants. **d)** Scatter plot of outgoing interaction strength (x-axis) and incoming interaction strength (y-axis) for AD participants. **e)** Heatmap of differential number of interactions. **f)** Heatmap of differential interaction strength. In the color bar, red represents an increase in signaling in the second dataset (AD) compared to the first (CT). **g)** Bar chart of AD versus CT pathways, showing the relative information flow. **h)** UMAP of AD and CT pathways grouped based on functional similarity, using the ‘uwot’ algorithm.

Analysis of the absolute numbers of interactions and strength measured for each group separately showed the Inflammatory microglia (Mic.15) as the dominant source of outgoing signaling in both groups, whereas Stress-response microglia (Mic.11) were the primary recipients of incoming signaling (**Figure 3c,d**). However, AD was characterized by stronger signaling through Mic.15 and Mic.11, whereas CT displayed a more balanced communication network across microglial populations. Differential changes were also observed for specific cell populations, including increased incoming and outgoing interactions in Mic.11 and reduced signaling in the Reactive cells of Mic.8 and the Proliferative Mic.1 in AD compared with CT, together with widespread changes in communication between individual cell-group pairs (**Figure 3e,f**).

To investigate coordinated communication programs, we applied NMF separately to each condition. In AD, outgoing signaling formed four patterns: Homeostatic-interferon signaling microglia, Stress-adapted lipid-regulatory microglia, Activated inflammatory and lipid-associated microglial output, and Peripheral monocyte communication (**Supplementary Figure S13**). Incoming signaling also formed four patterns: Homeostatic-reactive redox microglia, Peripheral monocyte communication, Stress-interferon responsive surveilling microglia, and Activated inflammatory and lipid-associated myeloid cells. In contrast, CT outgoing signaling was organized into six patterns: Microglia with homeostatic interferon signaling, Stress response, Microglia with homeostatic phagocytosis, Peripheral monocyte communication, Reactive-redox microglia due to lipid-associated inflammation, and Reactive-proliferative microglia, whereas incoming signaling formed four patterns: Homeostatic-reactive inflammatory redox interferon microglia, Peripheral monocyte communication, Stress-responsive proliferative surveilling microglia, and Activated myeloid cells and phagocytosis, associated with inflammation and lipids, indicating substantial reorganization of coordinated communication programs in AD (**Supplementary Figure S14** and **Supplementary Tables 11-14**).

Comparison of signaling pathway activity using paired Wilcoxon tests identified eight pathways with significantly higher activity in AD (ICAM, EDA, Resistin, Laminin, ADGRB, 12oxoLTB4, CD30, and JAM) and two pathways specific to CT (VISTA and CD200), while many pathways, including APP, Glutamate, Cholesterol, APOE, and SPP1, were active in both groups but differed in activity levels (**Figure 3g** and **Supplementary Figure S15**). Cell-type-specific pathway strength also differed between conditions: although Mic.15 remained the major signaling source, secretion strength was reduced across many pathways in AD, whereas Mic.11 showed enhanced incoming communication through selected pathways, including Glutamate and TGFβ (**Supplementary Figure S16**). Functional similarity analysis further grouped signaling pathways into four clusters, revealing distinct communication probability profiles between AD and CT, with Group 3 containing the largest number of functionally related pathways and showing prominent SPP1 activity in AD and CD45 activity in CT (**Figure 3h**).

The increased overall signaling observed in AD was accompanied by greater representation of specific subpopulations, while communication patterns were also reorganized across cell types and pathways.

### Persistent microglial signaling alterations beyond cell imbalance

To investigate the effects of cell imbalance, we repeated the cell-cell communication analysis between AD and CT using randomly sampled subsets matched for the number of participants in each group and the number of cells in each subpopulation, while preserving their relative abundance across glial populations. Several differences in specific cell-cell communication pairs were observed compared with the full analysis, indicating that the observed alterations in microglial communication were not primarily driven by differences in cell abundance between disease groups (**Figure 4**). The reduced signaling to Mic.8 (Reacting) in AD remained detectable after matching the number of participants and cells across glial subpopulations (**Figure 4a,b**). In addition, several communication differences involving the Reactive cells of Mic.6 became more pronounced after matching, with increased signaling from Mic.14 (Interferon response), Mic.15 (Inflammatory), and Mic.12 (Lipid-associated) in AD. These interactions were already mildly evident in the full analysis but became highly significant after accounting for differences in cell abundance. Conversely, the strong increase in signaling to and from the Stress-response cells of Mic.11 was no longer observed, suggesting that these interactions were primarily driven by the emergence of this stress-response cell state (**Figure 4c-f**).

**Fig 4.**
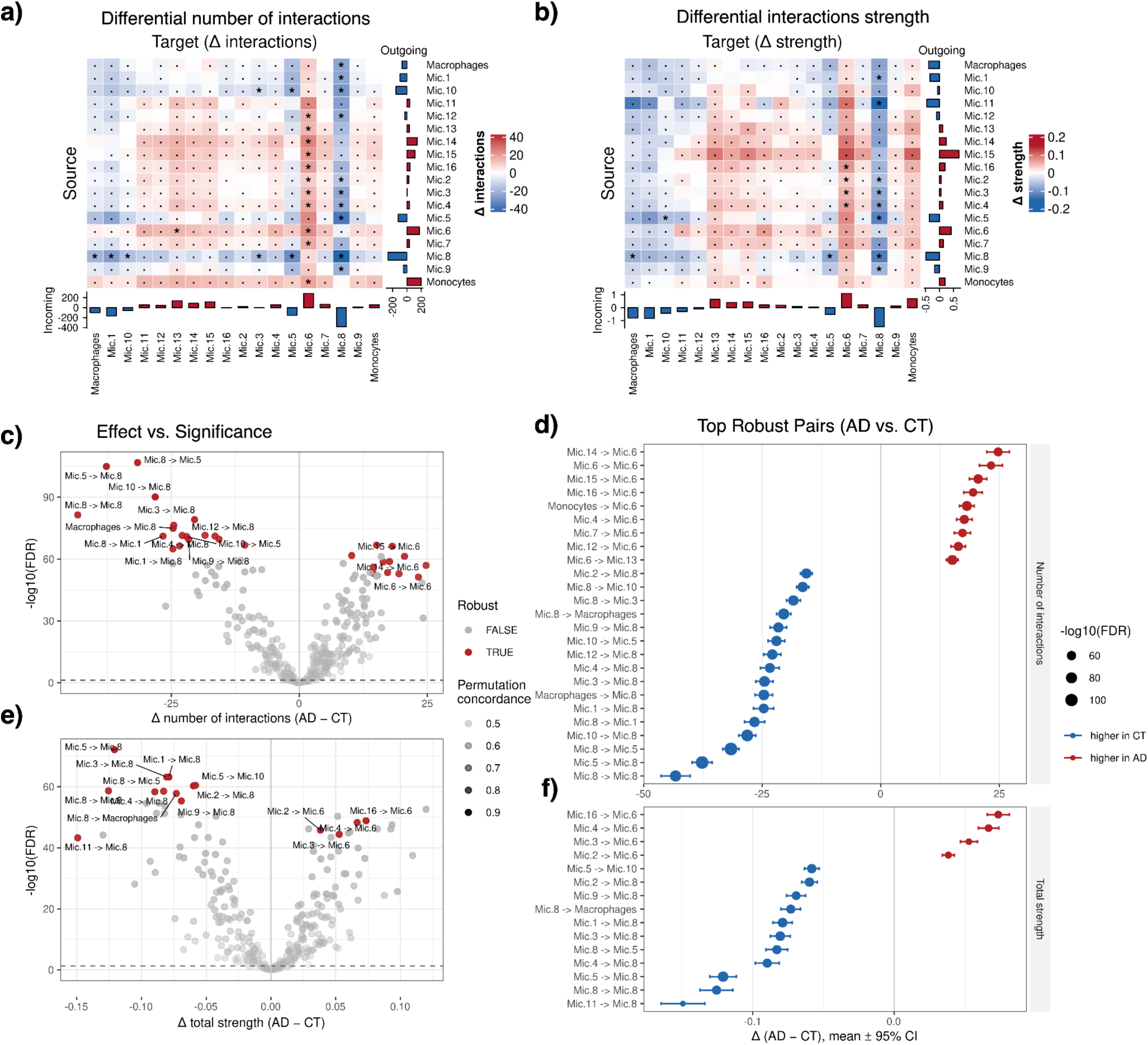
Differential cell–cell communication between AD and CT across matched-cell-count permutation runs. **a,b)** Heatmaps of Δ interaction count (a) Δ communication strength (b) between all source (rows) and target (columns) clusters. Red indicates higher signaling in AD, blue indicates higher signaling in CT. Top and side bar plots show each cluster’s total incoming and outgoing Δ, respectively. Asterisks (*) mark pairs that are robust (i.e. paired t-test FDR < 0.05 and concordance on direction across ≥ 90% of permutations); dots (·) mark pairs significant by FDR alone but with lower directional concordance across permutations. **c,e)** Volcano plots of Δ interaction count (c) and Δ communication strength (e) and versus statistical significance (-log_10_ FDR). Point color indicates robustness; point transparency reflects the fraction of permutations agreeing on effect direction. The top 15 robust pairs by |Δ| are labeled. **d,f)** Forest plots of the top 25 robust source → target pairs for each metric, faceted by Δ interaction count (e) and Δ communication strength (f). The dots represent the mean Δ (AD-CT) with 95% confidence intervals. Point color indicates the direction of the effect (red = higher in AD, blue = higher in CT); point size reflects −log_10_(FDR).

Overall, these results indicate that some alterations in microglial communication are robust to differences in cell abundance, whereas others could be largely attributed to changes in the cellular composition of the myeloid compartment.

### Associations of cell-cell interactions with cognitive decline and neuropathologies

To investigate the relationship between myeloid intercellular communication and AD-related phenotypes, we performed participant-level cell communication analyses and tested associations between personalized microglial cell-cell communication interaction strength versus clinical, pathological, vascular, and genetic features, including cognitive decline, cognitive resilience, PHFtau tangle density, amyloid-β load, global AD pathology, TDP-43, hippocampal sclerosis, vascular pathologies, *APOE* ε4 status, and AD diagnosis. In the discovery dataset, 24 microglial subpopulation pairs were significantly associated with AD-related phenotypes (adjusted *P*-value < 0.05), including 13 pairs associated with PHFtau density, 11 with amyloid-β load, 8 with global AD pathology, and 5 with hippocampal sclerosis. Consistent with previous reports identifying Mic.12 and Mic.13 as Lipid-associated microglial subpopulations implicated in AD proteinopathies (Green et al. 2024), the strongest association was observed between Mic.12 → Mic.13 signaling and PHFtau tangle density (-log_10_ *P*-value = 8.80), followed by the reciprocal interaction (-log_10_ *P*-value = 8.24). Interactions involving Mic.11, Mic.13, Mic.1 (Proliferative), and Mic.16 (SERPINE1^+^) as signaling sources were also strongly associated with PHFtau density but showed reduced strength for amyloid-β load. Mic.13 autocrine signaling showed the strongest association with amyloid-β load (-log_10_ *P*-value = 6.19). Additional associations highlighted specific communication programs, including Mic.16 outgoing signaling to multiple microglial populations associated with hippocampal sclerosis and Mic.12 autocrine signaling associated with cerebral amyloid angiopathy. Hierarchical clustering of significant communication pairs revealed coordinated modules, including closely related lipid-associated microglia (Mic.12 and Mic.13) and distinct signaling programs such as Mic.16 nuclear receptor signaling (**Figure 5a** and **Supplementary Tables 15-17**).

**Fig 5.**
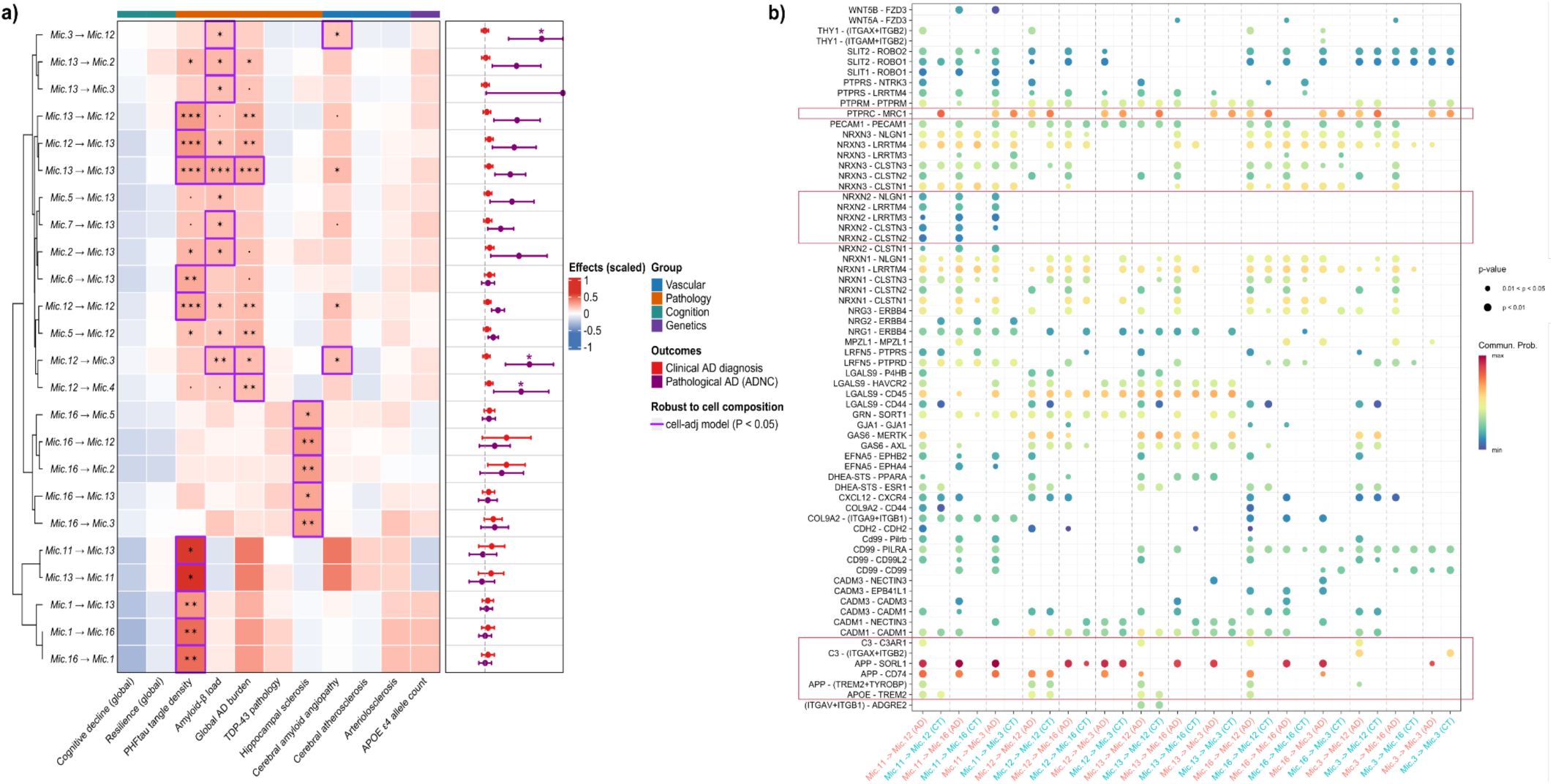
Association analysis with AD-related traits and genetics. **a)** Heatmap and forest plot showing the significant associations between sender → receiver cell group pairs and AD phenotypes. Associations of continuous and binary outcomes were measured via linear and logistic regression, respectively. In the heatmap, regression estimates are represented by color intensity and statistical significance showed as asterisks (FDR adjusted ***p<0.001, **p<0.01, *p<0.05). Phenotypes are labeled according to their category. Purple squares highlights results robust to cell composition adjustment. The forest plots show results for binary outcomes as log Odds Ratio (logOR) and 95% CI. Results nominally significant after cell composition adjustment are indicated with a purple asterisk. **b)** Bubble plot of cell communication signaling. Interactions are compared between CT and AD through the probability of communication. Pairs are read as L-R; the + sign within the parentheses indicates a multimeric receptor complex. Each interaction shown occurs between the emitter groups Mic.11, Mic.12, Mic.13, Mic.16, and Mic.3 and the receiver groups Mic.12, Mic.16, and Mic.3.

To evaluate if cell imbalance could affect the association between interaction strength and the AD-related phenotypes we included the cell counts relative to each cell pair in the regression models as covariates. Results were considered to be robust to cell composition if the original association was FDR < 0.05 and had a nominal *P*-value < 0.05 in the adjusted model. Many of the associations remained significant after controlling for the respective cell counts (**Figure 5a** and **Supplementary Figure 17a**). Importantly, the overall correlation was also highly correlated between models (Spearman *ρ* = 0.88) (**Supplementary Figure 17b**).

To further characterize the communication programs identified above, we examined the L-R interactions underlying the significant microglial cell-subpopulation pairs in AD and CT, highlighting how individual L-R pairs are differentially utilized depending on both disease status and the communicating cell populations. Overall, most interactions were increased or preferentially detected in AD, although differences depended on the communicating cell populations. Among the most prominent interactions were APP-SORL1, APP-CD74, and PTPRC-MRC1, which showed high communication probabilities in both conditions. However, many AD-enriched interactions involved Mic.11 as a signaling source, including an AD-specific NRXN2 ligand-associated interaction module. Additional interactions, including APP-SORL1, APP-CD74, APP-(TREM2+TYROBP), C3-C3AR1, and C3-(ITGAX+ITGB2), were preferentially observed in AD, whereas APOE-TREM2, PTPRC-MRC1, and glutamate pathway interactions were more similarly represented between AD and CT (**Figure 5b** and **Supplementary Figures S18-S19**).

Together, these findings demonstrate that microglial communication networks are extensively remodeled in AD, with disease-associated changes characterized by coordinated inflammatory and lipid-associated signaling programs linked to neuropathological accumulation.

### Replication in external datasets showed high validation rates

To validate the Discovery (CUIMC1) findings, we repeated the network inference and association analyses in three independent snRNA-seq datasets from the DLPFC: CUIMC2 (n=212), MIT (n=191), and Diversity (n=155), after removing individuals overlapping with the Discovery cohort. Although some participants were shared among replication datasets, analyses were performed separately by study to avoid repeated measures (**Figure 6a**). The cell communication analysis using all cells from each dataset identified 4,725 interactions in CUIMC2, 4,918 in MIT, and 4,799 in Diversity, representing 55, 46, and 42 inferred signaling pathways, respectively (**Supplementary Figures S20–S22** and **Supplementary Tables 18-20**). Across the Discovery and replication datasets, 38 signaling pathways were consistently detected, with 71 L-R pairs shared among the studies. While the Discovery dataset contained the largest number of unique interactions (93 exclusive L-R pairs), replication datasets showed greater overlap, supporting the reproducibility of the inferred myeloid communication network (**Figure 6b-c**).

**Fig 6.**
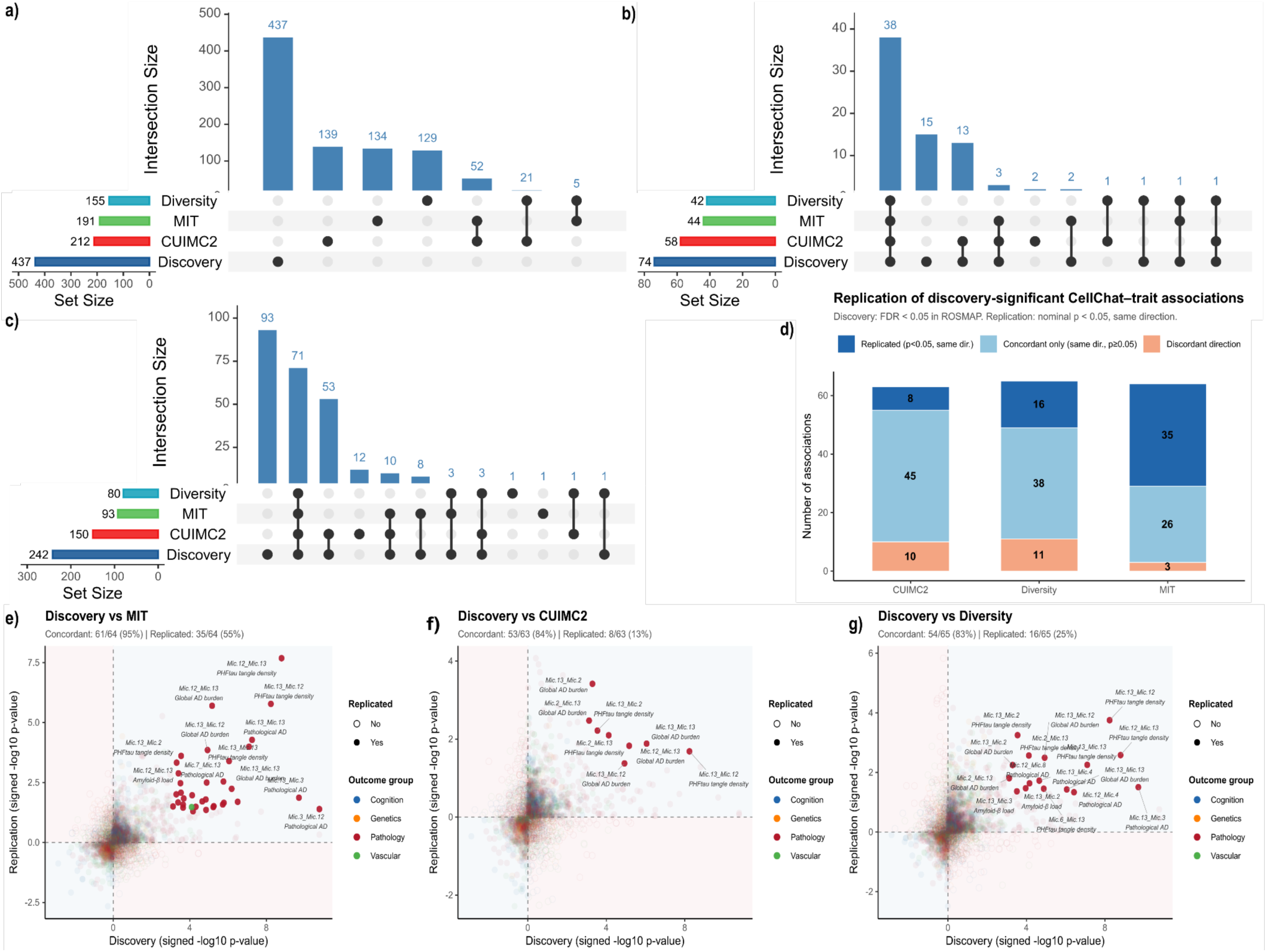
Replication analysis. **a)** Upset plot of the number of participants in the Discovery dataset (CUIMC1) and the replication datasets, as well as their intersections. **b)** Number of replicated pathways between the datasets, and **c)** Number of replicated L-R pairs. **d)** Only communications that were statistically significant in the Discovery data (FDR < 0.05) were included in the subsequent analysis. Dark blue indicates that the association reached a *P*-value < 0.05 in the replication dataset. Light blue indicates directional agreement, and orange indicates opposite direction. **e)** Representation of replication Discovery versus MIT. **f)** Discovery versus CUIMC2 and **g)** Discovery versus Diversity. The axes represent the transformed *P*-value (signed −log_10_ *P*-value), where the magnitude indicates the strength of the significance and the sign indicates the direction of the association. The blue quadrants indicate directional agreement, and the labeled points represent associations that reached statistical replication p < 0.05.

We next repeated participant-level cell communication association analyses across replication studies and combined all datasets, resulting in a total of 840 participants (**Supplementary Figure 23)**. Among 43 myeloid communication pairs tested, 42 were significantly associated with AD-related phenotypes, including PHFtau tangle density (28 pairs), amyloid-β load (24 pairs), global AD pathology (26 pairs), and hippocampal sclerosis (9 pairs). Importantly, most of these results were robust for cell composition adjustment (correlation of t-statistics Spearman *ρ* = 0.91; **Supplementary Figure 24**).

The strongest associations involved Mic.13 and Mic.12 interactions, particularly Mic.13 → Mic.12 (-log_10_ *P*-value = 15.20), Mic.12 → Mic.13 (-log_10_ *P*-value = 13.80), and Mic.13 → Mic.13 (-log_10_ *P*-value = 10.77), all associated with PHFtau tangle density. These interactions were also among the strongest associations with global AD pathology. Additional associations highlighted phenotype-specific communication programs, including Mic.15, Mic.16, Mic.12, and the Enhanced redox cells of Mic.9 as a signaling module associated with hippocampal sclerosis, Mic.13 → Mic.6 and Mic.13 → Mic.7 (Reacting, *TMEM163*+) interactions associated with cerebral atherosclerosis, and Mic.13 → Mic.4 (Surveilling) associated with *APOE* ε4 allele count. Categorical phenotype analyses further showed strong associations with pathological AD diagnosis, particularly involving Mic.13-related interactions (**Supplementary Table 21**).

To assess replication consistency between Discovery and external datasets, we evaluated concordance and replication rates across cohorts. MIT showed the highest replication performance, with 95% concordance, 55% replication, and only 4% discordant associations, including predominantly pathological associations and one vascular association. CUIMC2 and Diversity also showed high concordance (84% and 83%, respectively), with replication rates of 13% and 25% (**Figures 6d-g**). Overall, these results demonstrate that the identified myeloid communication associations are robust across independent datasets.

Importantly, these findings were replicated using distinct participant groups and snRNA-seq datasets generated by independent research groups. Furthermore, replication in the AMP-AD Diverse study, which includes individuals from non-European ancestries, supports the generalizability of these myeloid communication patterns across diverse populations.

## DISCUSSION

In this study, we investigated cell-cell communication among microglial states in the human DLPFC using snRNA-seq data from the ROSMAP cohorts. Through L-R network inference, we mapped signaling interactions among CNS myeloid populations and characterized the role of individual microglial subpopulations as signal senders and receivers. We found that AD pathology is associated with increased and more specialized microglial communication compared with cognitively unimpaired individuals. By integrating cell-communication metrics with neuropathological and clinical phenotypes, we identified specific cell-pair interactions associated with AD-related traits, many of which replicated across independent datasets. For example, lipid-associated microglia (Mic.12 and Mic.13) emerged as central hubs of signaling associated with tau pathology, whereas nuclear receptor signaling microglia (Mic.16) showed strong associations with hippocampal sclerosis. These findings suggest that distinct microglial states contribute differentially to specific pathological features of AD.

The identification of functionally distinct microglial states has substantially advanced our understanding of neurodegenerative diseases. Consistent with previous studies describing disease-associated microglial phenotypes, including MGnD, ARM, HAM, and LDAM states (Krasemann et al. 2017; Sala Frigerio et al. 2019; Marschallinger et al. 2020), our results indicate that microglial subpopulations undergo transcriptional and functional specialization in the AD brain. The communication patterns inferred from our analysis support the concept of microglial reprogramming during neurodegeneration and extend prior observations by revealing how these states coordinate through distinct signaling pathways. We also observed that Inflammatory microglia (Mic.15) dominate outgoing communication, whereas Stress-associated microglia (Mic.11) preferentially receive signals, suggesting complementary roles in propagating and sensing pathological stimuli. Furthermore, the increased communication activity observed in AD relative to non-cognitive impaired participants corroborates previous reports of enhanced L-R signaling in AD brains (Zhang et al. 2023).

Our findings provide insights into the cellular mechanisms through which microglial states may contribute to AD pathology. The central role of lipid-associated microglia in tau-related signaling networks suggests that these cells may participate in coordinating responses to neurofibrillary tangle accumulation. In contrast, the association between Mic.16 signaling and hippocampal sclerosis points to potentially distinct microglial programs involved in non-AD neurodegenerative pathology. We also identified signaling pathways uniquely present in AD, including the ICAM and JAM adhesion pathways. The expression of ICAM signaling by inflammatory microglia is particularly interesting given prior evidence linking ICAM1 to plaque-associated neuroinflammation and microglial chemotaxis in AD (Akiyama et al. 1993; Janelidze et al. 2018; Wang et al. 2024). Together, these results indicate that AD is characterized not only by increased microglial communication but also by the emergence of specialized signaling programs that may contribute to disease progression.

An important consideration in comparing cell-cell communication between AD and CT is that differences in cellular composition can influence inferred communication networks. To address this potential confounding effect, we performed a cell abundance–matched permutation analysis that simultaneously matched the number of participants and cells across microglial subpopulations. This analysis revealed that several communication changes persisted after accounting for differences in cell abundance, supporting the presence of disease-associated alterations in microglial signaling that are not simply a consequence of changes in cell composition. In particular, reduced signaling involving the Reacting Mic.8 population remained robust in AD, while interactions involving the Reactive Mic.6 population, including signaling from Interferon-response Mic.14, Inflammatory Mic.15, and Lipid-associated Mic.12, became more pronounced after matching. Conversely, the increased signaling involving Stress-response Mic.11 was no longer detected after accounting for cell abundance, suggesting that this alteration is largely attributable to the expansion of this cell state in AD, which is also biologically relevant. Together, these findings highlight the importance of considering cellular composition when interpreting cell-cell communication networks and distinguish communication alterations that reflect changes in microglial abundance from those that represent persistent rewiring of intercellular signaling.

A strength of this study is the use of single nucleus human brain datasets, enabling high-resolution characterization of microglial communication networks in the aging and AD cortex. The combination of cell-communication inference, neuropathological associations, and replication across independent cohorts provides robust evidence for the biological relevance of the identified interactions. In addition, our analyses generate a comprehensive catalog of candidate L-R interactions that can serve as a foundation for future mechanistic studies. Several limitations should also be considered. The ROSMAP cohorts are enriched for highly educated individuals and predominantly represent people of European ancestry, potentially limiting generalizability, although we could replicate some of our findings in a Diverse cohort. Also, snRNA-seq captures only the nuclear transcriptome and miss signaling components primarily represented in the cytoplasm. Finally, L-R inference relies on currently available databases (CellChat DB in this case), which remain incomplete and may not capture all biologically relevant interactions. Although our analyses provide a comprehensive view of microglial communication in the aging and AD cortex, future work is needed to experimentally validate these predicted interactions and determine how they influence disease progression and therapeutic response.

## METHODS

### Study participants

Data were obtained from participants of two longitudinal clinical-pathological cohort studies: the Religious Orders Study (ROS) and the Rush Memory and Aging Project (MAP), collectively referred to as ROSMAP. Both studies were approved by an Institutional Review Board of Rush University Medical Center (Chicago, IL). The ROS was established in 1994 and includes priests, brothers, and nuns from across the United States, while the MAP began in 1997 and includes individuals from retirement communities and private residences in northeastern Illinois (Bennett et al. 2018). At the time of enrollment, all participants were without known dementia and provided informed consent for annual clinical and cognitive assessments, and an Anatomic Gift Act for brain donation after death. The phenotypic variables included in this manuscript are described below.

### Demographic covariates Age at death

Age of death is calculated from subtracting date of birth from date of death and dividing the difference by days per year (365.25).

### Years of education

The years of education variable is based on the number of years of formal schooling reported at baseline cognitive testing (Bennett et al. 2005).

### Sex

Sex is self-reported.

### Genetics

DNA genotyping was performed on peripheral blood mononuclear cells or brain tissue using high-throughput sequencing at codons 112 (position 3,937) and 158 (position 4,075) of exon 4 of the *APOE* gene (Yu et al. 2017).

### Clinical and neuropathological evaluations

During neuropathological evaluations, examiners were blinded to participants’ clinical data to ensure unbiased assessments. To determine the probability of AD pathology, a dichotomized version of the National Institute on Aging - Alzheimer’s Association (NIA-AA) criteria was used by a neuropathologist to classify the likelihood of a pathological AD diagnosis into two levels: intermediate-to-high probability or low probability, based on the presence of neurofibrillary tangles (Braak) and neuritic plaques (CERAD) (D. A. Bennett et al. 2006).

Brains were removed, weighed, and fixed in 4% paraformaldehyde. Tissue samples were collected from eight brain regions: angular gyrus, anterior cingulate, calcarine cortex, entorhinal cortex, hippocampus, inferior temporal cortex, mid-frontal gyrus, and superior frontal cortex. To quantify amyloid-β deposition and PHFtau tangle density, slides were prepared from 20-micrometer sections for immunohistochemical staining with specific antibodies. Whole-slide images of the eight regions were scanned using software to perform the counts (Bennett et al. 2003; Kapasi et al. 2023).

### AD-Related Dementia (ADRD) pathology

#### Amyloid-β measurements

To measure beta-amyloid load, 20-micron sections are prepared from eight brain regions: the angular gyrus, anterior cingulate, calcarine cortex, entorhinal cortex, hippocampus, inferior temporal gyrus, middle frontal gyrus, and superior frontal gyrus. Histochemical and digital analyses were performed to delineate regions of interest and quantify them into digital scores for each individual slide; these scores represent the percentage of tissue occupied by beta-amyloid following a square-root transformation (Kapasi et al. 2023).

#### Tangles density

A similar process is used to measure tangle density; however, in this case, the total tangle density for each individual slide is calculated by summing the nuclei with medium and strong thresholds and dividing by the area of analysis, resulting in a value of Phftau positive tanglecounts per mm², followed by the application of a square-root transformation (Kapasi et al. 2023).

#### AD pathology and other neurodegenerative diseases

The global AD pathological burden summarizes quantitative measures of neuritic plaques, diffuse plaques, and neurofibrillary tangles in silver-stained slides across five brain regions: the prefrontal cortex, pre-temporal cortex, inferior parietal cortex, entorhinal cortex, and hippocampus, examined via microscopy (Bennett et al. 2003). Other evaluated pathologies included TDP-43 staining, classified as present or absent in the amygdala, or extending to other regions (Nag et al. 2018). Additionally, hippocampal sclerosis was graded from 0 to 5, being considered severe at stage 5 in the CA1 or subiculum (Nag et al. 2015).

#### Vascular pathologies

Cerebral infarcts were defined as macroscopic and microscopic by standardized neuropathological examination (Schneider et al. 2005; Arvanitakis et al. 2011). Arteriosclerosis is assessed in small blood vessels using histological slides, while cerebral atherosclerosis is assessed in macroscopic arteries, both being semiquantitatively classified from 0 to 3 (Buchman et al. 2011; Arvanitakis et al. 2017). Cerebral amyloid angiopathy (CAA) evaluates the meningeal and parenchymal vessels across four regions and is also classified semiquantitatively (0–3) (Love et al. 2014; Boyle et al. 2015).

#### Cognitive decline and resilience

Cognitive function in ROSMAP participants was assessed through a battery of 21 performance tests, administered annually by examiners blinded to previous results. To construct a composite measure of global cognition, scores from 17 of the 21 tests were used, standardized into z-scores using the baseline mean and standard deviation, and then averaged to derive a composite global cognition measure (Wilson et al. 2015). The annual rate of change for each individual was estimated using a linear mixed-effects model with global cognition as the longitudinal outcome, adjusted for age at baseline, sex, and years of education (De Jager et al. 2012). To calculate cognitive resilience, the same model was applied to measure the slope of global cognition, controlling for the same demographic and adding neuropathological variables, including global AD pathology, amyloid-β, neurofibrillary tangles, Lewy body disease, TDP-43 staining, infarcts, hippocampal sclerosis, and vascular pathologies (Boyle et al. 2021).

The clinical diagnosis of cognitive status is performed by analyzing the results of the 21 tests. A neurologist with expertise in dementia reviews the clinical results without prior knowledge of age or any autopsy data (Schneider et al. 2007; David A. Bennett et al. 2006). A diagnosis is made for the most likely clinical diagnosis at the time of death, classified as Alzheimer’s dementia versus mild cognitive impairment (MCI) or no cognitive impairment (NCI) (David A. Bennett et al. 2006; Bennett et al. 2002).

#### Single-nucleus RNASeq dataset (Discovery - CUIMC1)

Expression data were generated at the Columbia University Irving Medical Center (CUIMC) from samples provided by the Rush Alzheimer’s Disease Center of the ROSMAP participants. Specifically, in the study by Green et al. (2024), snRNA-seq was performed on postmortem DLPFC. The samples were received frozen and contained varying amounts of gray matter, white matter, and adherent meninges. While still frozen, the white matter and meninges were carefully removed during dissection.

The frozen gray matter was then subjected to lysis and mechanical homogenization, followed by centrifugation with samples processed two at a time. This step separated the nuclei from the cytoplasm, resulting in nuclear pellets that were kept on ice while the supernatant was discarded. The nuclei were resuspended in a PBS solution containing an RNase inhibitor, filtered, and counted. Nuclear profiling was conducted on pooled batches of 40,000 nuclei, comprising 5,000 nuclei from each of 8 participants, with the exception of batch B63, which contained 7 participants. Sample assignment to batches was randomized to mitigate batch effects, and was balanced for clinical and pathological diagnosis, as well as sex. These pools of 40,000 nuclei were then processed on the 10x Genomics platform for snRNA-seq library construction, following the manufacturer’s protocol. The resulting libraries were sequenced at the Broad Institute using Illumina HiSeqX and at the New York Genome Center using Illumina NovaSeq 6000, and the data from both platforms were combined for the final analysis (Green et al. 2024).

#### Data for replication

##### CUIMC2 dataset

In the study by (Comandante-Lou et al. 2025), data were generated at the Columbia University Irving Medical Center (CUIMC) from samples provided by the Rush Alzheimer’s Disease Center, originating from 240 individuals in the Religious Orders Study (ROS) and the Rush Memory and Aging Project (MAP) cohorts. The snRNA-seq was performed using tissue samples from the DLPFC. The tissue samples underwent mechanical homogenization and centrifugation. The suspension was then filtered and underwent density gradient centrifugation; this step separated the nuclei from the cytoplasm, resulting in nuclear pellets that were kept on ice while the supernatant was discarded. The pooled nuclei were then centrifuged, resuspended, filtered, and counted at 21,500 nuclei. With the nuclei isolated, the 10x multiome protocol (Chromium Next GEM Single Cell Multiome ATAC + Gene Expression Reagent Bundle) was followed according to the manufacturer’s instructions.

After the GEMs were generated, libraries were prepared and sequenced on an Illumina NovaSeq 6000 system. Next, the sequences were aligned to the hg38 human genome using CellRanger (v2.0.2) and the CellBender tool to remove ambient RNA technical noise. Finally, to identify which cell belonged to which study sample, demultiplexing was performed using the demuxlet (v1.11) software. The data were then normalized and underwent quality control, so that the cells could be annotated through reference mapping.

##### MIT dataset

Data were generated by the Massachusetts Institute of Technology (MIT) of samples originating from 427 individuals in the Religious Orders Study (ROS) and the Rush Memory and Aging Project (MAP) cohorts, provided by the Rush Alzheimer’s Disease Center. The snRNA-seq was performed using frozen tissue samples from the DLPFC, following the protocol described by Mathys et al. (2023).

Samples were stored on ice at 4 °C and homogenized using a Wheaton Dounce tissue homogenizer and filtered. Nuclei were separated from the cytoplasm by centrifugation, resulting in nuclear pellets that were kept on ice while the supernatant was discarded. The isolated nuclei were then counted and diluted to a concentration of 1,000 nuclei per microliter. Library construction was performed using the 10x Genomics Chromium platform (v2 and v3). Finally, the libraries were sequenced on Illumina machines (HiSeq and NovaSeq) (Mathys et al. 2023).

##### Diversity dataset

Data were generated by the Accelerating Medicines Partnership for Alzheimer’s Disease Diversity (AMP-AD Diversity) project from samples of individuals enrolled in the Religious Orders Study (ROS), Rush Memory and Aging Project (MAP), Minority Aging Research Study (MARS), African American Clinical Core (AA-Core), and Latino Core Study (LATIC), provided by the Rush Alzheimer’s Disease Center (Luquez et al. 2026). The data used in this study were derived from samples of participants with diverse genetic backgrounds, including non-Hispanic White, Latino, and African American individuals. During the DLPFC dissection, the white matter and meninges were carefully removed while the tissue remained frozen in blocks. The frozen gray matter then underwent lysis and mechanical homogenization, followed by centrifugation in a swinging-bucket rotor, with samples processed in pairs. Nuclei separated from the cytoplasm were kept as pellets on ice, while the supernatant was discarded and the remaining tissues were processed to complete a batch of 8 samples. The nuclei were resuspended in a PBS solution containing an RNase inhibitor, filtered, and counted. Nuclear profiling was performed with approximately 20,000 nuclei from 3 samples, comprising different donor/region combinations pooled into a single sample, following the 10x multiome protocol (Chromium Next GEM Single Cell Multiome ATAC + Gene Expression Reagent Bundle) according to the manufacturer’s instructions. After GEM generation, libraries were prepared and sequenced on an Illumina NovaSeq 6000 system at the New York Genome Center. Subsequently, reads were aligned to the human genome using CellRangerArc (v2.0.2) and the CellBender tool to remove background noise. Finally, demultiplexing was performed using the Freemuxlet workflow from the popscle package, run on each 10x Genomics run, to identify which study sample each cell belonged to.

Finally, the cell annotations were anchored to the Discovery dataset so that we could ensure that Mic.1, for example, has the same signature across all Mic.1 from the four datasets used. The myeloid cells (microglia, infiltrating monocytes, and macrophages) were selected for downstream analysis in this manuscript.

### Seurat pipeline

As detailed by Green et al. (2024), the raw RNA count matrix was processed using Seurat, an open-source R package for single-cell omics analysis (Hao et al. 2021, 2024). The analytic pipeline, executed with Seurat v.4.1.0, began with data normalization and scaling, followed by dimensionality reduction via Principal Component Analysis (PCA) and cell clustering.

A rigorous quality control step was then performed, applying cell-type-specific thresholds to remove low-quality nuclei with a low number of UMIs or unique genes, as well as doublets. Subsequently, cells were automatically annotated using an ElasticNet-regularized logistic regression classifier trained on a previously published cell atlas. This classified all nuclei into eight major cell types: excitatory neurons, inhibitory neurons, astrocytes, microglia, oligodendrocytes, OPCs, endothelial cells, and pericytes. For data visualization, a unified Uniform Manifold Approximation and Projection (UMAP) space was generated. Due to the large number of nuclei, a reference UMAP was created from a data subset, and the remaining nuclei were projected onto this space. Following this global clustering, a second round of clustering (subclustering) was performed separately on each cell type to identify distinct cell subpopulations.

Differentially expressed genes (DEGs) between clusters were identified using a negative binomial test, controlling for confounding variables. Finally, functional annotation of these DEGs was performed to identify enriched biological pathways (KEGG, Reactome) and Gene Ontology (GO) terms, thereby revealing the biological functions associated with each cell cluster. Microglia were annotated in 16 subpopulations, and the respective data were subjected to the “sctransform” method.

### Cell-cell communication analysis and NMF clusterization

In our study, we used the standard CellChat (v2.1.2) workflow using snRNA-seq data as input (Jin et al. 2021, 2025). CellChat is a tool that quantifies and analyzes cell-cell communication networks using single-cell transcriptomic data. Communication inference is based on the CellChatDB database, which is manually curated and supported by literature. This database accounts for the multimeric structure of L-R complexes, as well as soluble and membrane-bound stimulatory and inhibitory cofactors, such as agonists, antagonists, and co-receptors. L-R pairs are categorized into secreted signaling, extracellular matrix-receptor interaction, cell-cell contact, and non-protein signaling, and are further classified into functionally related signaling pathways. To quantify the probability of communication between cell groups, the algorithm employs a simplified model based on the law of mass action. Each interaction receives a score calculated from the average expression values of a ligand in the sender cell, its cognate receptor in the target cell, and their respective cofactors. To infer the myeloid communication network, we used 84,064 cells of the DLPFC from 437 ROSMAP participants, which were previously annotated by Green et al. (2024) into 16 subtypes of microglia, macrophages, and infiltrating monocytes (18 groups) using Seurat. Two analyses were performed: (1) including cells from all participants and (2) comparing AD cases and controls, characterizing our Discovery set. We created the CellChat object from the “SCT” assay. Using the respective L-R pair database (CellChatDB), we performed cell-cell communication network inference using the “truncatedMean” and “trim = 0.1” parameters for both the all-cell analysis and for each condition (comparison of AD versus CT cells).

Analysis 1 focused on building a cell communication map of myeloid cells in the aging brain. First, we presented the dominant sender and receiver cell groups in the inferred network using the standard CellChat function “netAnalysis_signalingRole_scatter,” which performs a centrality analysis using weighted directed network measures. Next, we highlighted the results of latent pattern inference with cell groups, performed using the non-negative matrix factorization (NMF) method based on Cophenetic and Silhouette metrics, utilizing a stabilized number of patterns. To visualize the identified outgoing and incoming patterns, we used a cutoff of 0.4 in the “netAnalysis_river” function.

Analysis 2 focused on highlighting differences found in brains with and without AD pathology; for this, we filtered cells by the AD pathological diagnosis variable. This analysis included 53,938 cells from 274 individuals with AD and 30,126 cells from 163 individuals in the control group, and was performed following the CellChat (v2) comparison pipeline. Comparative analysis is conducted through manifold learning and by crossing communication probabilities with differential gene expression analyses. Starting with the overall network signaling in each group, we highlighted the difference in the number and total strength of inferred interactions in each group (“compareInteractions”). We also compared the role of each microglial group side-by-side across both conditions. The differential number and strength of interactions (“netVisual_heatmap”) were calculated by subtracting the communication probability matrices quantitatively inferred for each group based on L-R pair expression.

To understand signaling pathway activity, we performed pathway clustering based on functional similarity across the two groups (“netVisual_embeddingPairwise”); this analysis was conducted through signaling network classification learning. We again applied the NMF method to compare the inferred outgoing and incoming communication patterns in each condition. Regarding pathway activity in AD and CT, we performed relative information flow per pathway, calculated using the “rankNet” function, applying the Wilcoxon test and a “cutoff = 0.05”. Finally, we visualized the dysfunctional signaling of L-R interactions in the cell groups of interest across both sets, comparing communication probabilities with the standard “netVisual_bubble” function. Differential expression (DE) analysis was performed first by identifying genes that changed significantly based on fold change, followed by extracting L-R pairs where ligands were up-regulated and calculating an enrichment score for each.

### Cell abundance–matched CellChat analysis

To mitigate potential effects of cell imbalance, we repeated the cell communication analysis using random subsets matched for both participant number and cell abundance between AD and CT groups. A total of 293 permutations were performed, each restricting the analysis to 100 AD and 100 CT participants. Within each permutation, cells were randomly sampled to match the number of cells in each glial subpopulation between groups while preserving the relative abundance of each subpopulation within the overall pooled composition. Subpopulations containing fewer than 10 cells in either group were excluded from the corresponding permutation.

CellChat was independently rerun on each matched subset, yielding distributions of interaction counts and communication strengths for each source-target cluster pair across the 293 permutations. The same parametrization of the full analysis was used. For each pair and metric, the paired difference (AD - CT) was calculated within each permutation. The mean difference across permutations was tested against zero using a paired t-test. *P*-values were corrected for multiple comparisons across cluster pairs using the Benjamini–Hochberg procedure. For each pair, we also calculated the 95% confidence interval of the mean difference and the concordance rate, defined as the proportion of permutations in which the direction of the AD - CT difference matched the direction of the overall mean difference. Pairs were considered robust if they showed FDR-adjusted significance (q < 0.05) and ≥ 90% concordance across permutations.

### Association analysis

To evaluate the association between microglial intercellular communication and AD-related traits, we initially executed the CellChat pipeline individually for each participant (n=437), using the same parameter as before. We obtained a total of 428 CellChat objects; some participants did not have sufficient cells (<10 per subtype) for robust cellular network inference and were excluded. This step generated an aggregated interaction strength (weight) matrix representing the communication intensity of all microglia pairs (sender → receiver) per person.

Associations were evaluated using a linear regression model for continuous variables (Cognition, Pathology, Vascular, and Genetics groups) and logistic regression for categorical variables (Clinical AD Diagnosis and Pathological AD). All models were adjusted for age, sex, and years of education to control for potential confounding factors. For continuous phenotypes, the strength of association was assessed using *P*-values and the regression coefficient (β), outcomes were standardized before regressions and in the heatmap visualization colors were scaled to the maximum absolute β across outcomes. For categorical phenotypes, associations were quantified by Odds Ratios (OR) and Z-scores, with 95% confidence intervals (CI) presented on a logarithmic scale. All tests were adjusted using the Benjamini-Hochberg method to control for the false positive rate (FDR). Finally, we applied hierarchical clustering to group cell pairs with similar associative L-R communications, other details are described in the figures legends.

### Replication analysis with external datasets

To replicate the discovery findings, we utilized three external datasets described by Comandante-Lou et al. (2025): MIT (Massachusetts Institute of Technology); CUIMC2 (Columbia University Irving Medical Center 2); and Diversity (Accelerating Medicines Partnership for Alzheimer’s Disease Diversity project). Participants from CUIMC2 and MIT are part of ROSMAP, while participants in the Diversity project are from ROSMAP as well as the Minority Aging Research Study (MARS), the African American Clinical Core, and the Latino CORE Study (LATC). This includes individuals who are African American (69%), White Hispanic (25%), or American Indian or Alaska Native (5%), based on self-reported race. All five longitudinal studies are conducted at Rush University, and all participants underwent the same ante-and post-mortem procedures. Consistent with the data used in the Discovery analysis, the data used in the replication analysis are derived from brain samples of the DLPFC.

For a valid replication, we removed participants overlapping with the Discovery dataset, totaling 191 individuals from MIT, 212 from CUIMC2, and 155 from Diversity, for a total of 558 participants. To ensure consistency of cell states across different datasets, reference mapping was performed using the Seurat package, where the cell states previously described by Green et al. (2024) (CUIMC1-Discovery) were used as a reference atlas for the three datasets (CUIMC2, MIT, and Diversity), which were treated as queries. We repeated the association analysis between intercellular communication and AD phenotypes to investigate whether the findings from the Discovery phase would be replicated using external data. Following the same steps, we inferred the communication network using the previously described parameters individually for each participant across the three groups. We performed the association analysis between cell pairs and AD-related traits using a linear regression model for continuous variables and a logistic regression model for categorical variables, with the same parameters as the Discovery phase, adjusted for age, sex, and years of education. To allow for direct comparison between different models, the logistic regression coefficients (Odds Ratios) were log-transformed (log(OR)). The results were presented in a heatmap with combined associations from the replication and discovery phases, totaling 840 participants.

For the comparison across cohorts, we aligned the results using a unique association key linking the predictor cell pair to the outcome phenotype. We calculated the −log_10_(p)×sign(β) metric to summarize the statistical significance strength and the direction of the association’s effect. As replication criteria, we defined that an interaction is only considered replicated if it achieves an FDR ≤ 0.05 in the discovery phase, shows the same direction of effect (concordance), and has a nominal p < 0.05 in the replication cohort. Interactions that maintained the direction of effect but did not reach nominal significance were classified only as concordant. Finally, discordant interactions were defined by the reversal of the coefficient sign between cohorts.

## SUPPLEMENTAL INFORMATION

Supplementary Figures 1-24 in the PDF file.

Supplementary Tables 1-21 in Excel format.

## ACKNOWLEDGMENTS

We thank all the RADC cohort participants and the investigators, and the staff at the Rush Alzheimer’s Disease Center. The five RADC cohorts are supported by P30AG10161, P30AG72975, R01AG15819, R01AG17917, U01AG46152, U01AG61356, and R01AG22018. The data available in the AD Knowledge Portal would not be possible without the participation of research volunteers and the contribution of data by collaborating researchers. The results published here are in whole or in part based on data obtained from the AD Knowledge Portal (https://adknowledgeportal.org). Study data were provided by the Rush Alzheimer’s Disease Center, Rush University Medical Center, Chicago, IL, USA. Data collection was supported through funding by NIA grants P30AG10161, P30AG72975, R01AG17917, R01AG015819 (ROS and MAP), RF1AG057473 (CUIMC1, single-nucleus RNA-seq), R01AG074003 and UO1NS110453 (MIT, single-nucleus RNA-seq), R01AG067025, U01AG061356 and R01AG066831 (Diverse, single-nucleus RNA-seq), U01AG061356, U01AG072572 and R01AG036042 (CUIMC2, single-nucleus RNA-seq).

## AUTHOR CONTRIBUTIONS

L.S performed data analysis, including running the CellChat pipeline, NMF clusterization, and AD vs CT analysis; wrote the manuscript with K.P.L. Next, R.A.V wrote the functions for association analysis with the AD-related traits and performed the permutation analysis across cell subpopulations. N.C.L mapped the cell type annotations for replication analysis. G.G clusterized and annotated the Discovery dataset. S.T and Y.W provided feedback. N.H supervised the clusterization and annotation of the Discovery dataset. V.M supervised the clusterization and annotation of the Diversity dataset. P.L.D.J provided the snRNASeq; supervised the clusterization and annotation of the Discovery dataset. R.T.R supervised the cell-communication analysis with K.P.L. Also, D.A.B built the ROSMAP cohorts and much of the omics; made critical comments on the manuscript. K.P.L conceived, supervised this study and wrote the manuscript with the co-authors feedback.

## DATA AVAILABILITY

Transcriptomic data used in this manuscript are available with the following accession numbers: Discovery CUIMC1 (Synapse ID: syn31512863), CUIMC2 (Synapse ID: syn71711894), Accelerating Medicines Partnership for AD Diversity project (Synapse ID: syn52339332), MIT (Synapse ID: syn52293417). The phenotype data can be requested at the RADC Resource Sharing Hub at www.radc.rush.edu or the AD Knowledge Portal adknowledgeportal.synapse.org. The relevant code generated by this work is available on the GitHub repository https://github.com/RushAlz/Mic_cell_comm_public_release/.

