## Supplementary figures for "Altered microglial communication in brains with Alzheimer’s Disease pathology"

### List of Supplementary Figures

**Session 01: All myeloid cells.** **3**
Supplementary Figure 1: The ligand–receptor (L-R) interaction database. 3
Supplementary Figure 2: Signaling sent by each group of cells. 4
Supplementary Figure 3: Heatmap of the number of interactions. 5
Supplementary Figure 4: Heatmap of interaction strength. 6
Supplementary Figure 5: Incoming versus outgoing microglia signaling. 7
Supplementary Figure 6: Inference of the number of outgoing patterns. 8
Supplementary Figure 7: Visualization of the identified outgoing patterns. 9
Supplementary Figure 8: Inference of the number of incoming patterns. 10
Supplementary Figure 9: Visualization of the identified incoming patterns. 11
Supplementary Figure 10: Heatmap of the strength of the outgoing and incoming interactions. 12
Supplementary Figure 11: Signaling groups with similar functionalities. 13
**Session 02: CT vs AD.** **14**
Supplementary Figure 12: Differences in microglial cell-cell communication between AD and CT. 14
Supplementary Figure 13: Outgoing patterns in CT vs AD. 15
Supplementary Figure 14: Outgoing signalling patterns CT vs AD. 16
Supplementary Figure 15: Relative information flow via CT and AD. 17
Supplementary Figure 16: Incoming signaling patterns CT vs AD. 18
**Session 03: Replication analysis.** **19**
Supplementary Figure 17: Replication of association analysis with AD phenotypes. 19
Supplementary Figure 18: Dysfunctional signaling in CT vs AD. 20
Supplementary Figure 19: Word cloud. 21
Supplementary Figure 20: Signaling pathways of the CUIMC2. 22
Supplementary Figure 21: Signaling pathways of the Diversity dataset. 23
Supplementary Figure 22: Signaling pathways of the MIT. 24
Supplementary Figure 23: Association analysis with AD-related traits in datasets combined. 25
Supplementary Figure 24: Comparison of AD-trait association models adjusted and not adjusted by cell counts using combined datasets. 26

**List of Supplementary Tables (Separate spreadsheets)**

Supplementary Table 1: Phenotype data for the Discovery dataset.
Supplementary Table 2: Brief description of myeloid subpopulation of cells.
Supplementary Table 3: Cell communication results for all cells.
Supplementary Table 4: 2,425 variable ligand-receptor (L-R) pairs.
Supplementary Table 5: 242 unique L-Rs.
Supplementary Table 6: Number of detected cells and genes by cell type.
Supplementary Table 7: Non-negative matrix factorization (NMF) results for the outgoing comparison using all cells.
Supplementary Table 8: NMF results for the incoming comparison using all cells.
Supplementary Table 9: Cell communication results for the cells from participants with AD pathology.
Supplementary Table 10: Cell communication results for the cells from participants without AD pathology (CT).
Supplementary Table 11: NMF results for the outgoing comparison for the AD participants.
Supplementary Table 12: NMF results for the incoming comparison for the AD participants.
Supplementary Table 13: NMF results for the outgoing comparison for the CT participants.
Supplementary Table 14: NMF results for the incoming comparison for the CT participants.
Supplementary Table 15: Association results for the Discovery dataset (n = 437). Complete table.
Supplementary Table 16: Values represent point estimates results of regression models. Beta for linear regression, and ORs for logistic regression.
Supplementary Table 17: Association results for the Discovery dataset. Table with the p-values from the regression analysis.
Supplementary Table 18: Cell communication results for the CUIMC2 microglia dataset.
Supplementary Table 19: Cell communication results for the MIT microglia dataset.
Supplementary Table 20: Cell communication results for the AMP-AD Diversity microglia dataset.
Supplementary Table 21: Association results for the combined dataset (n = 840). Complete table.

**Session 01: All myeloid cells.**

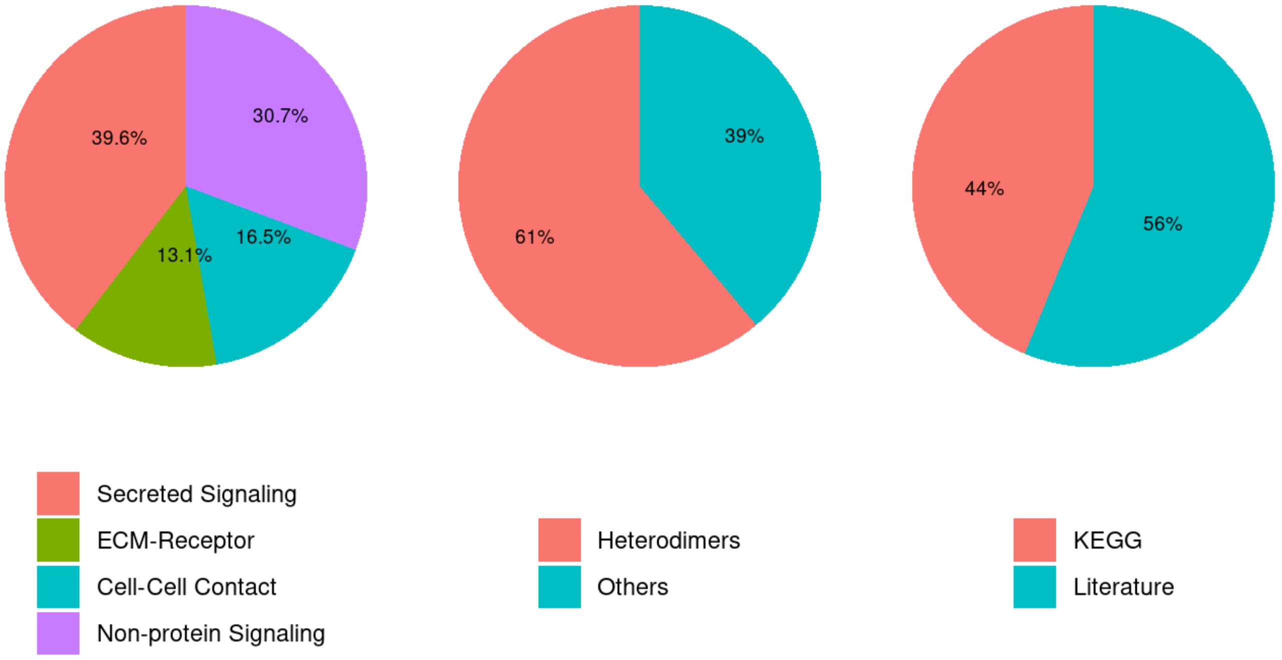


**Supplementary Figure 1: The ligand–receptor (L-R) interaction database.**
CellChatDB is a manually curated database of literature-supported ligand–receptor interactions, containing 3,234 validated molecular interactions. These interactions are categorized into different types, as shown in the pie charts: "Secreted Signaling," "ECM–Receptor," "Cell–Cell Contact," and "Non-protein Signaling." By default, "Non-protein Signaling" interactions are not used.

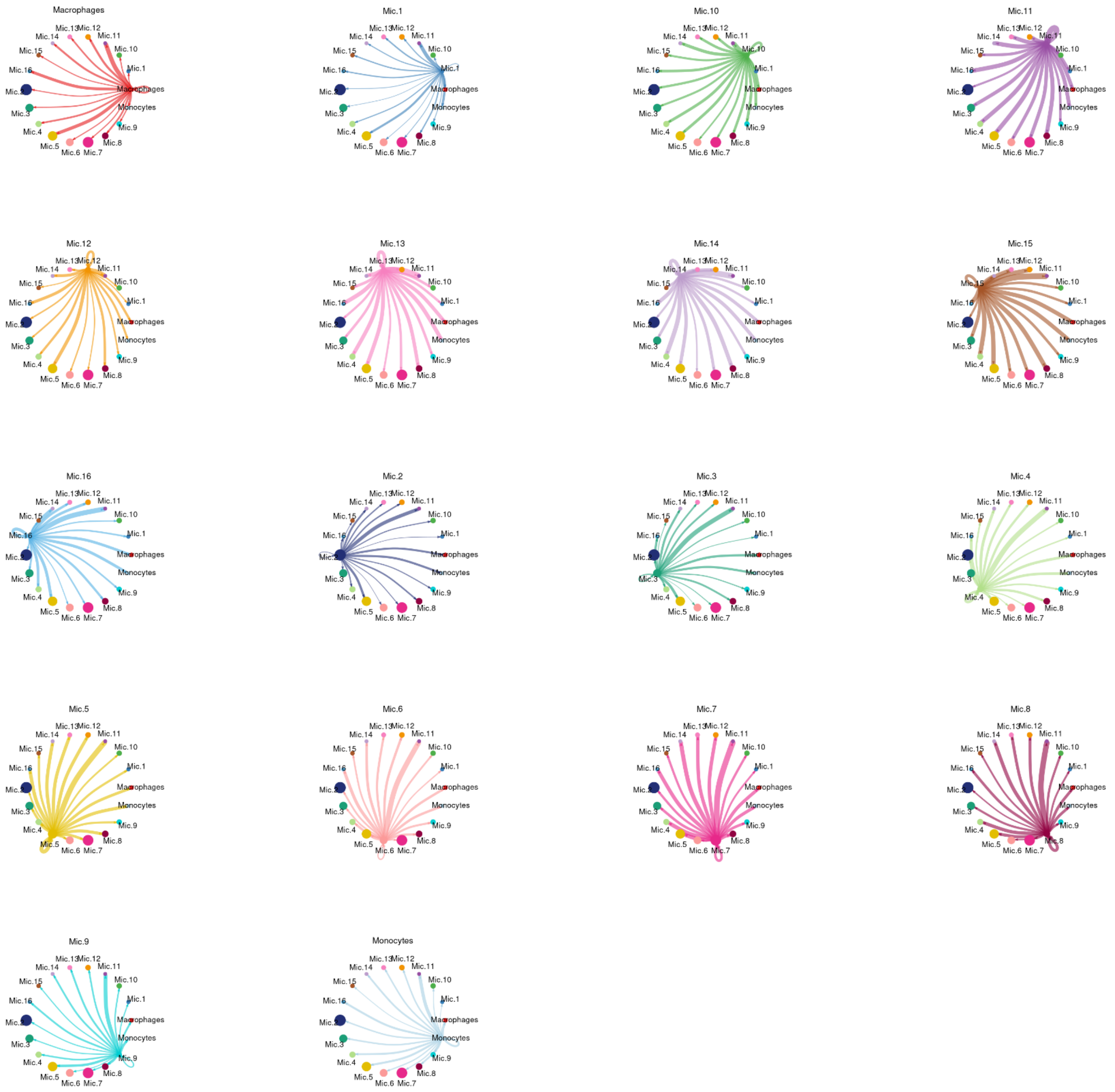


**Supplementary Figure 2: Signaling sent by each group of cells.**
The width of the edges represents the strength of the communication.

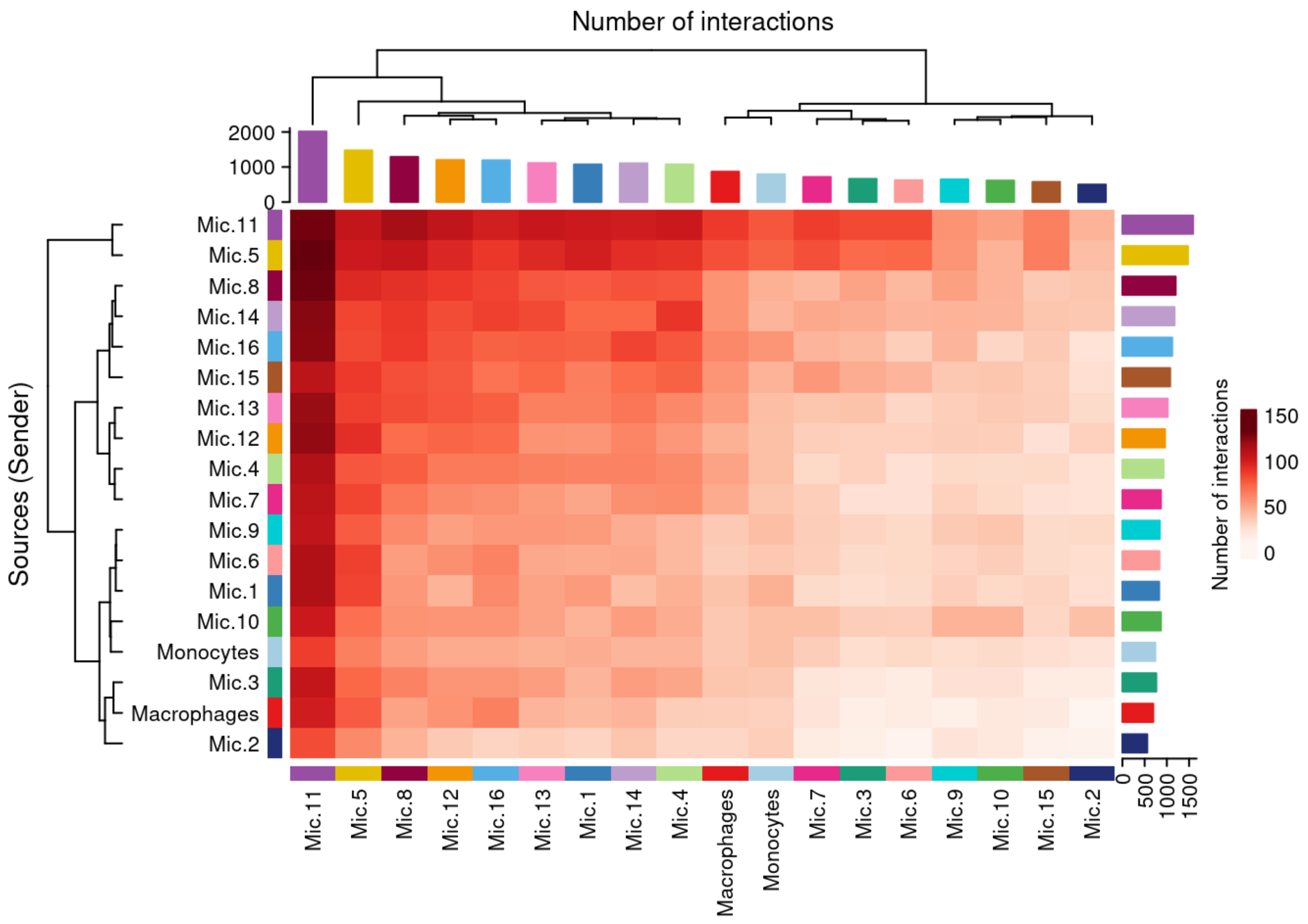


**Supplementary Figure 3: Heatmap of the number of interactions.**
The intensity of the colors indicates a greater number of interactions between a sending cell group (Y-axis) and a receiving cell group (X-axis).

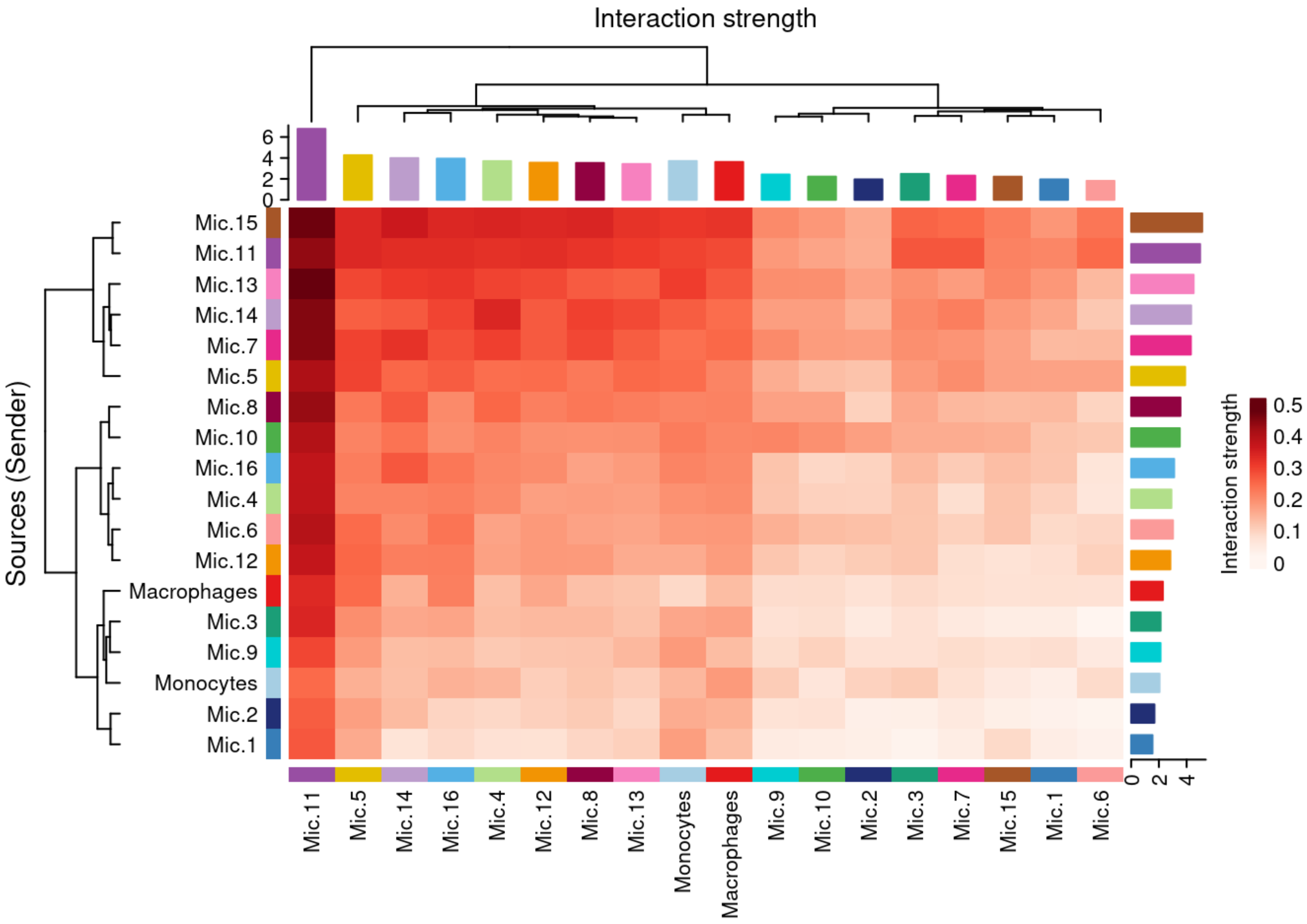


**Supplementary Figure 4: Heatmap of interaction strength.**
The intensity of the colors indicates greater interaction strength between a sending cell group (Y-axis) and a receiving cell group (X-axis).

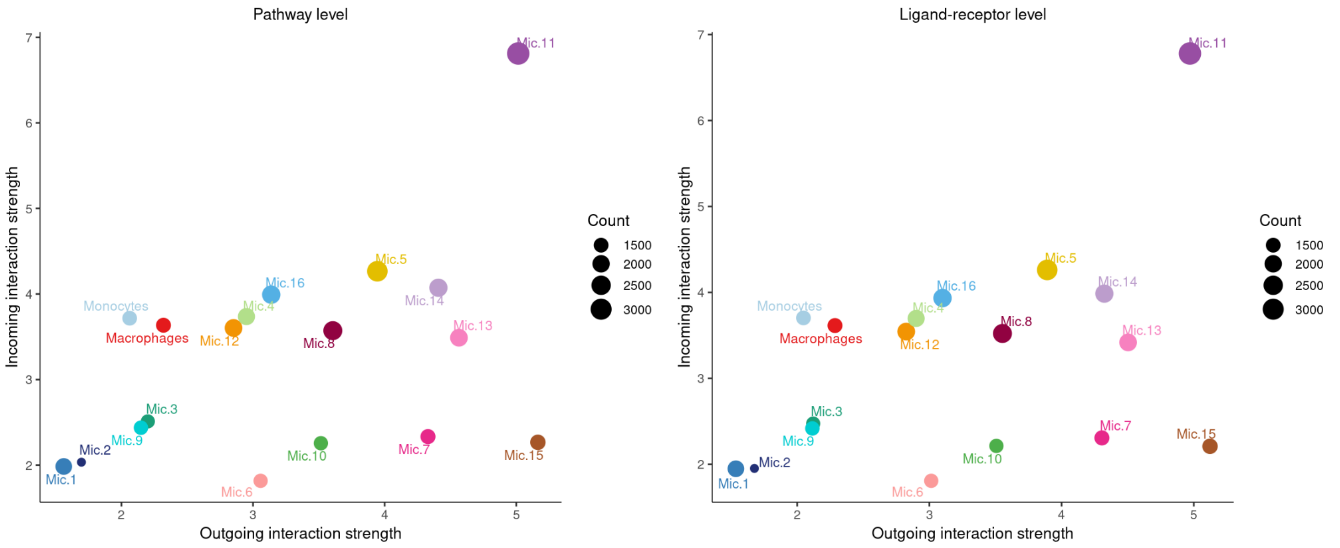


**Supplementary Figure 5: Incoming versus outgoing microglia signaling.**
In this scatter plot, the cell groups furthest to the right (X-axis) are sending more signals, while the groups further up (Y-axis) are receiving more signals. The left panel shows the role of cell groups at the pathway level, and the right panel at the ligand-receptor level.

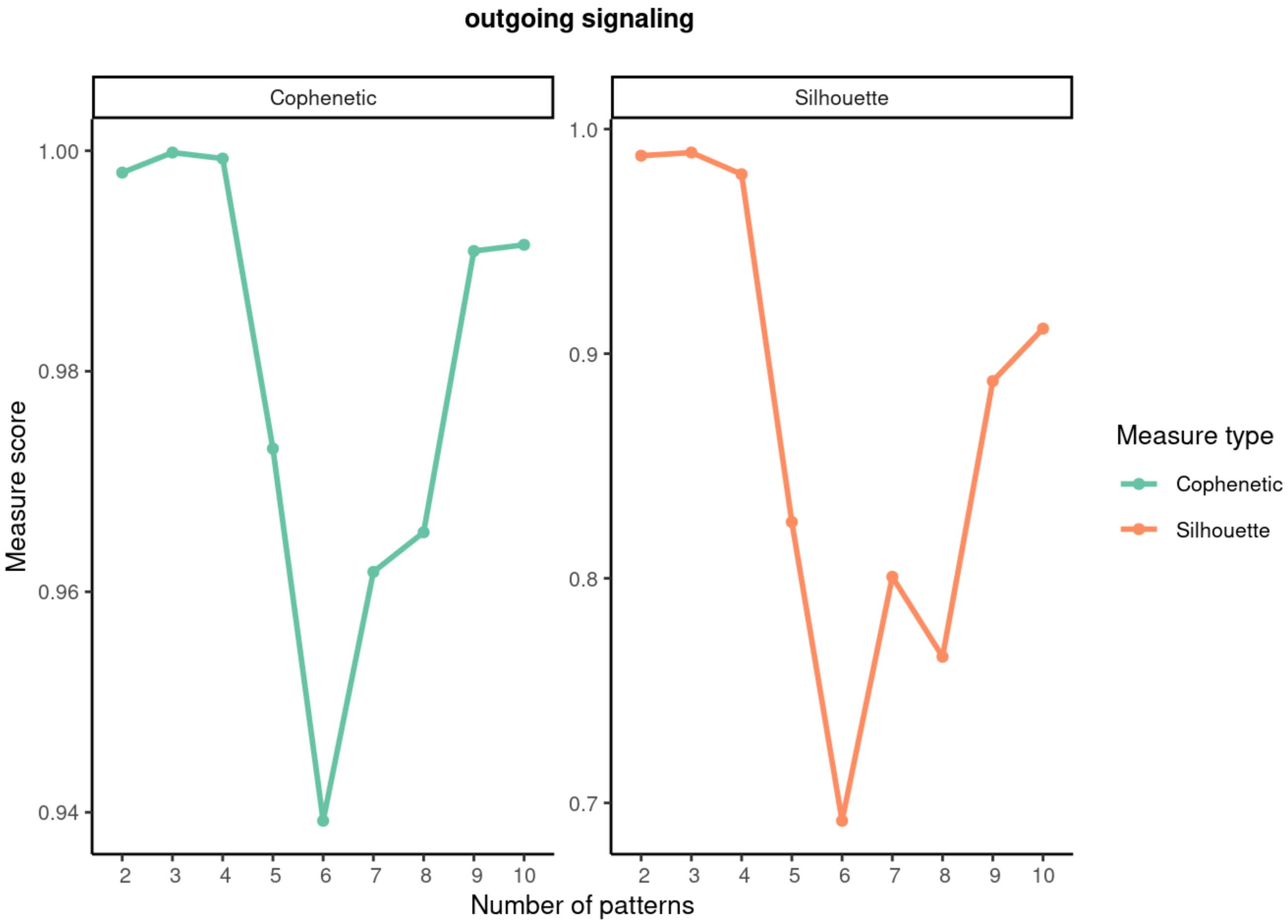


**Supplementary Figure 6: Inference of the number of outgoing patterns.**
Outgoing patterns show how the sender cells (signal source) coordinate with each other and with the signaling pathways to drive communication. The appropriate number of patterns is that in which the cophenetic and silhouette values begin to drop abruptly.

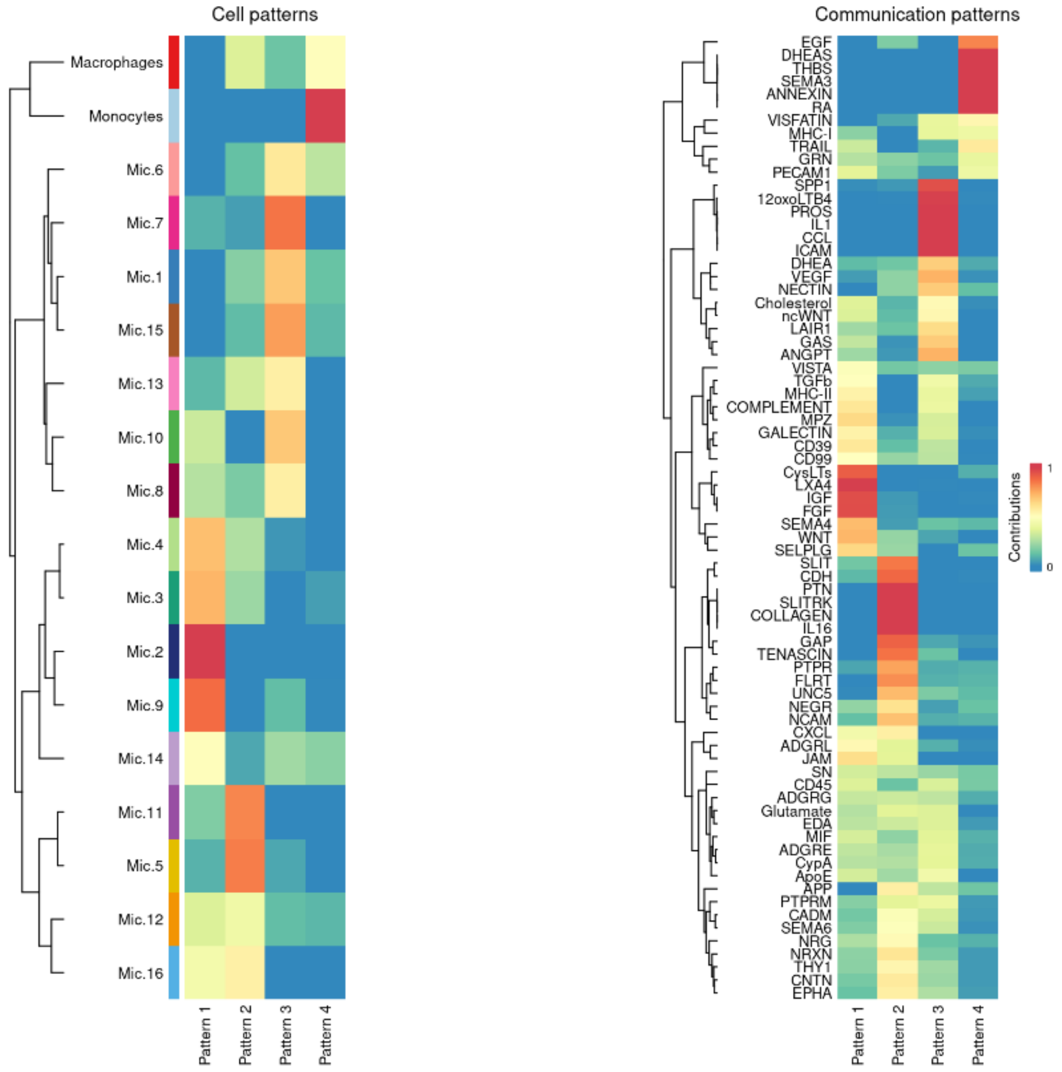


**Supplementary Figure 7: Visualization of the identified outgoing patterns.**
Heatmap displaying the most enriched cell groups and signaling pathways associated with each inferred pattern.

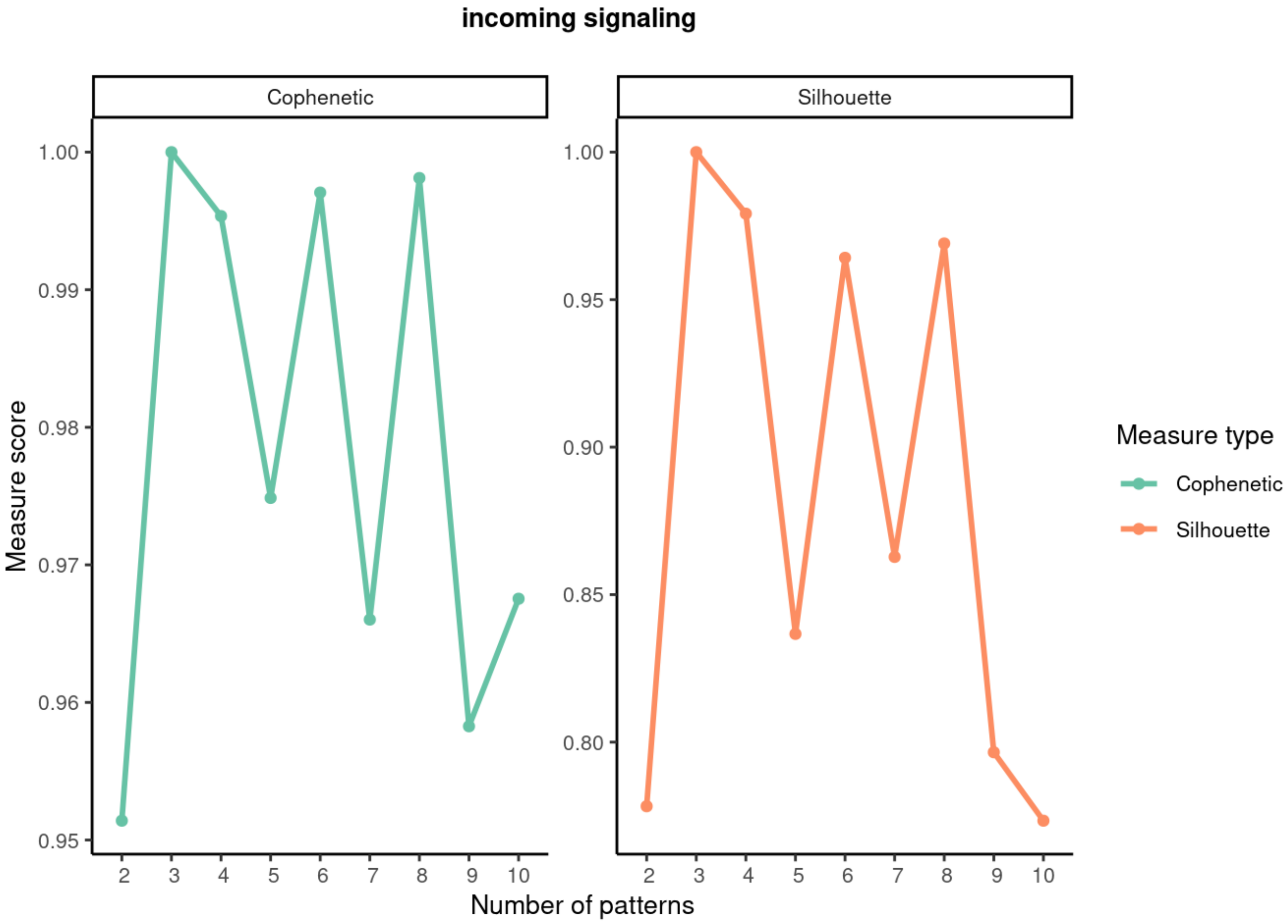


**Supplementary Figure 8: Inference of the number of incoming patterns.**
Incoming patterns show how target cells (signal receptors) coordinate with each other and with signaling pathways to respond to received signals. The appropriate number of patterns is that at which cophenetic and silhouette values begin to drop abruptly.

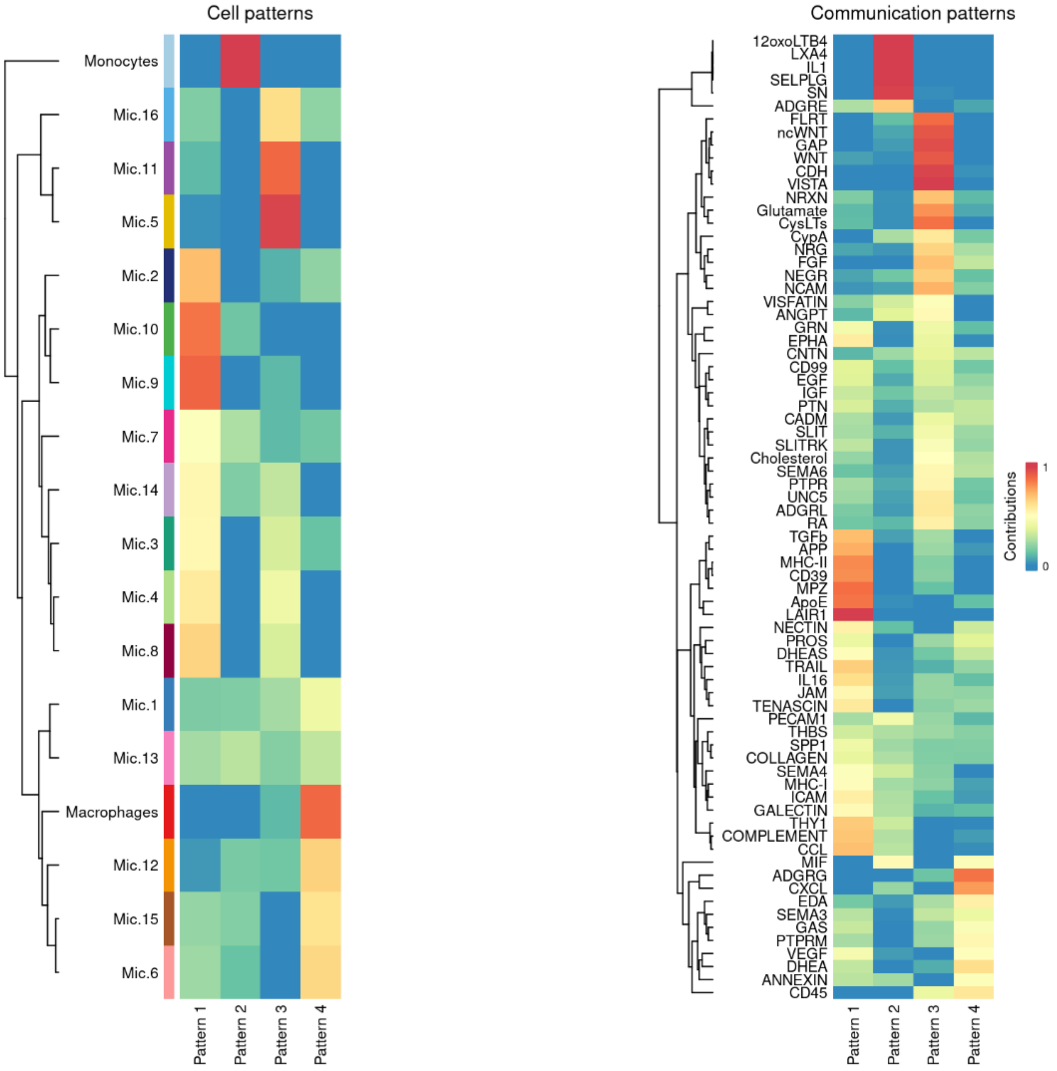


**Supplementary Figure 9: Visualization of the identified incoming patterns.**
Heatmap displaying the most enriched cell groups and signaling pathways associated with each inferred pattern.

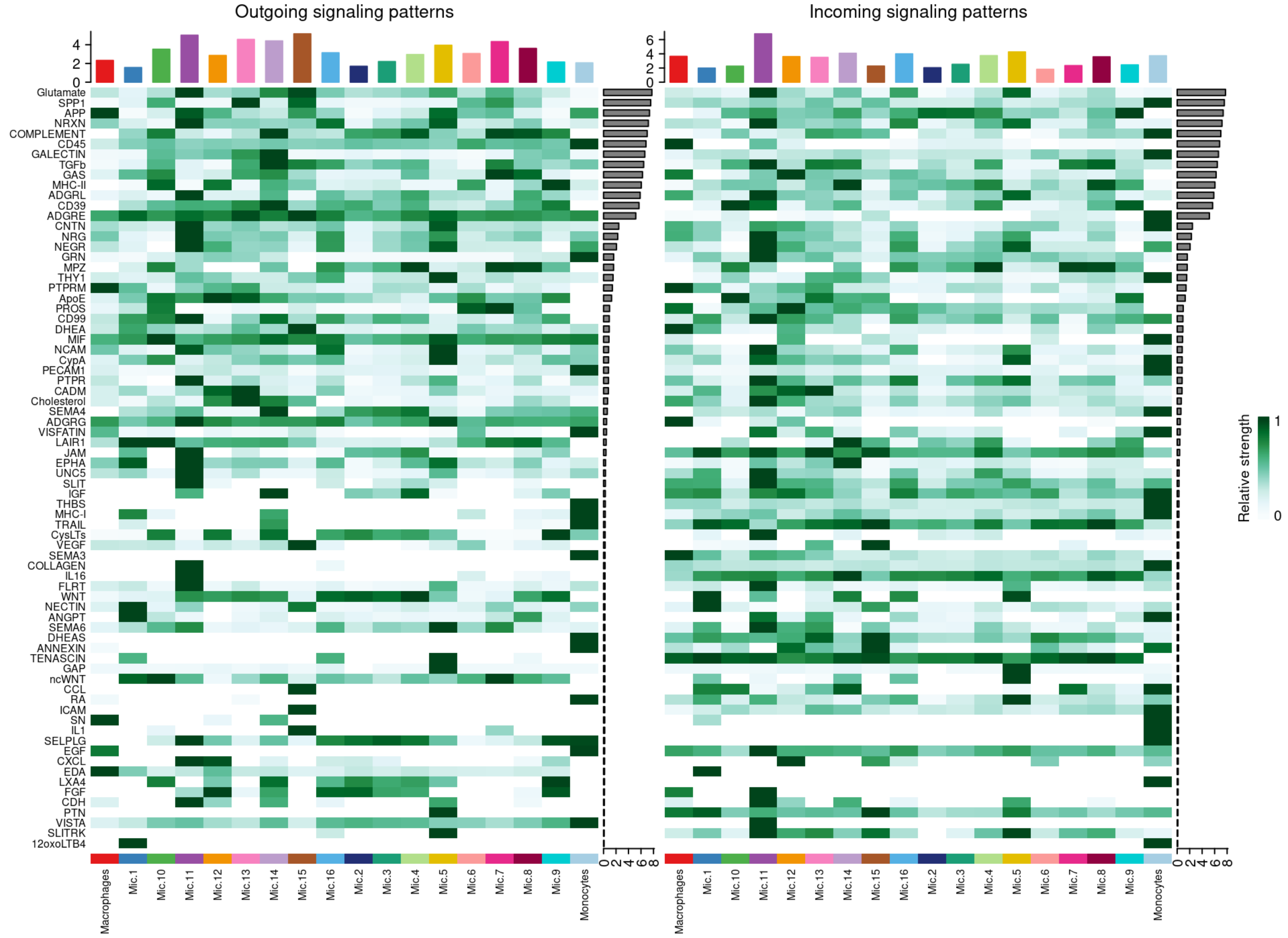


**Supplementary Figure 10: Heatmap of the strength of the outgoing and incoming interactions.** The intensity of the colors indicates the strength of the outgoing (left panel) and incoming (right panel) interactions of the signaling pathways in each group of cells.

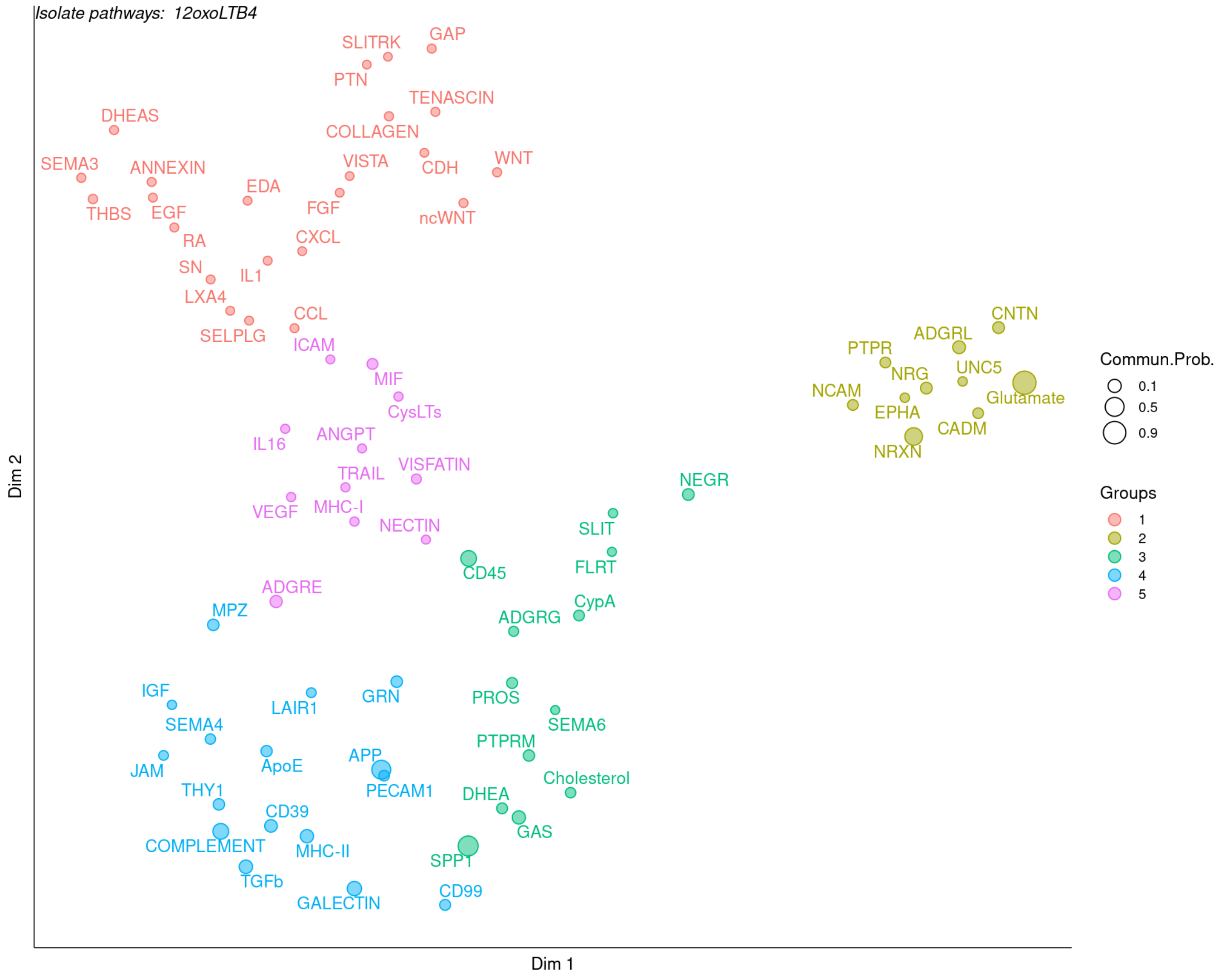


**Supplementary Figure 11: Signaling groups with similar functionalities.**

The grouping of signaling routes based on similarity and probability of communication can be visualized in a 2D space.

**Session 02: CT vs AD.**

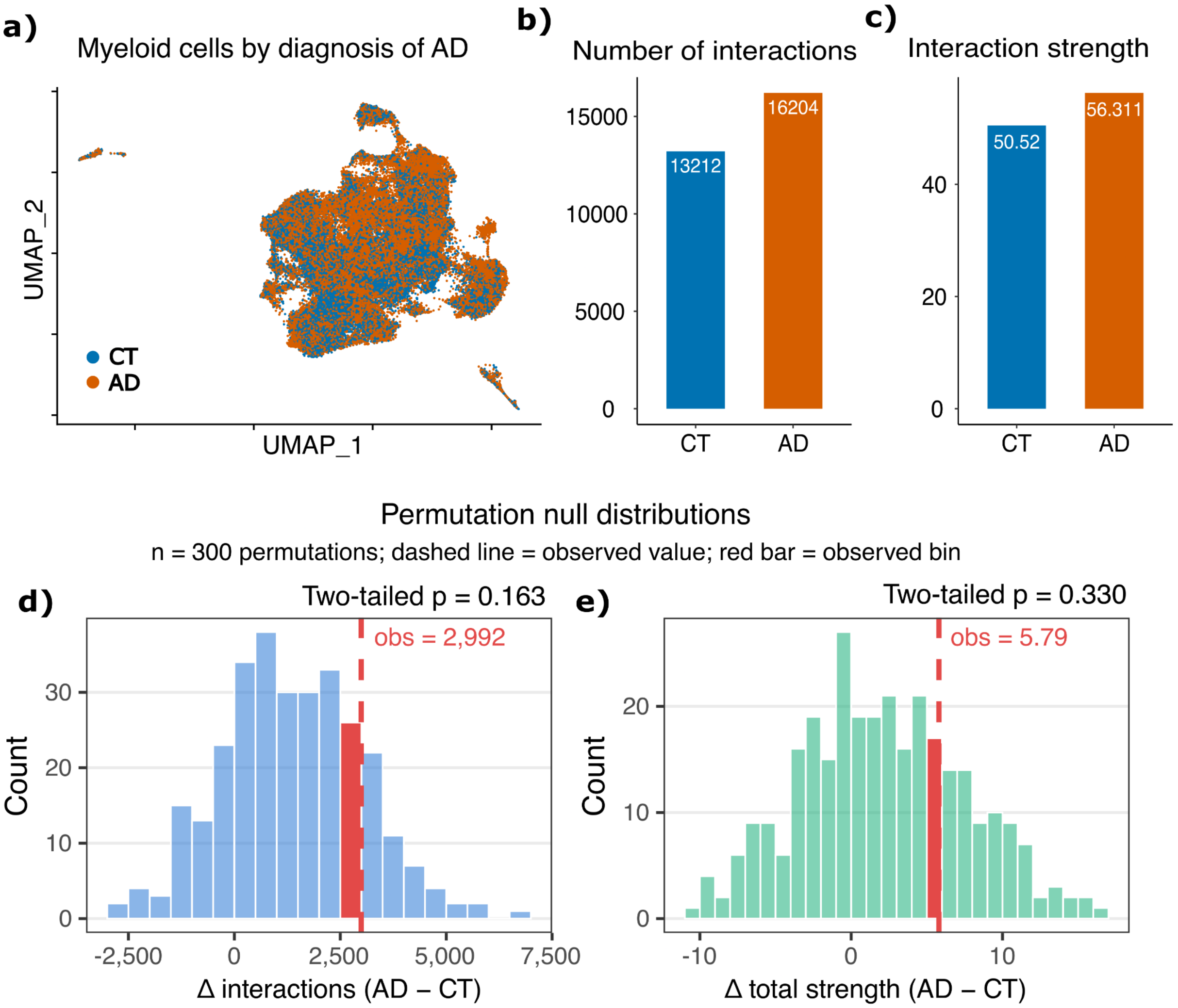


**Supplementary Figure 12: Differences in microglial cell-cell communication between AD and CT.**
**a)** UMAP colored by pathological diagnosis. **b-c)** Bar chart showing the number of interactions **(b)** and their strength **(c)**. **d)** Histogram of the permutation null distribution for the difference in the number of interactions between AD and CT. **e)** Histogram of the permutation null distribution for the difference in total interaction strength between AD and CT, based on cell–cell communication weights.

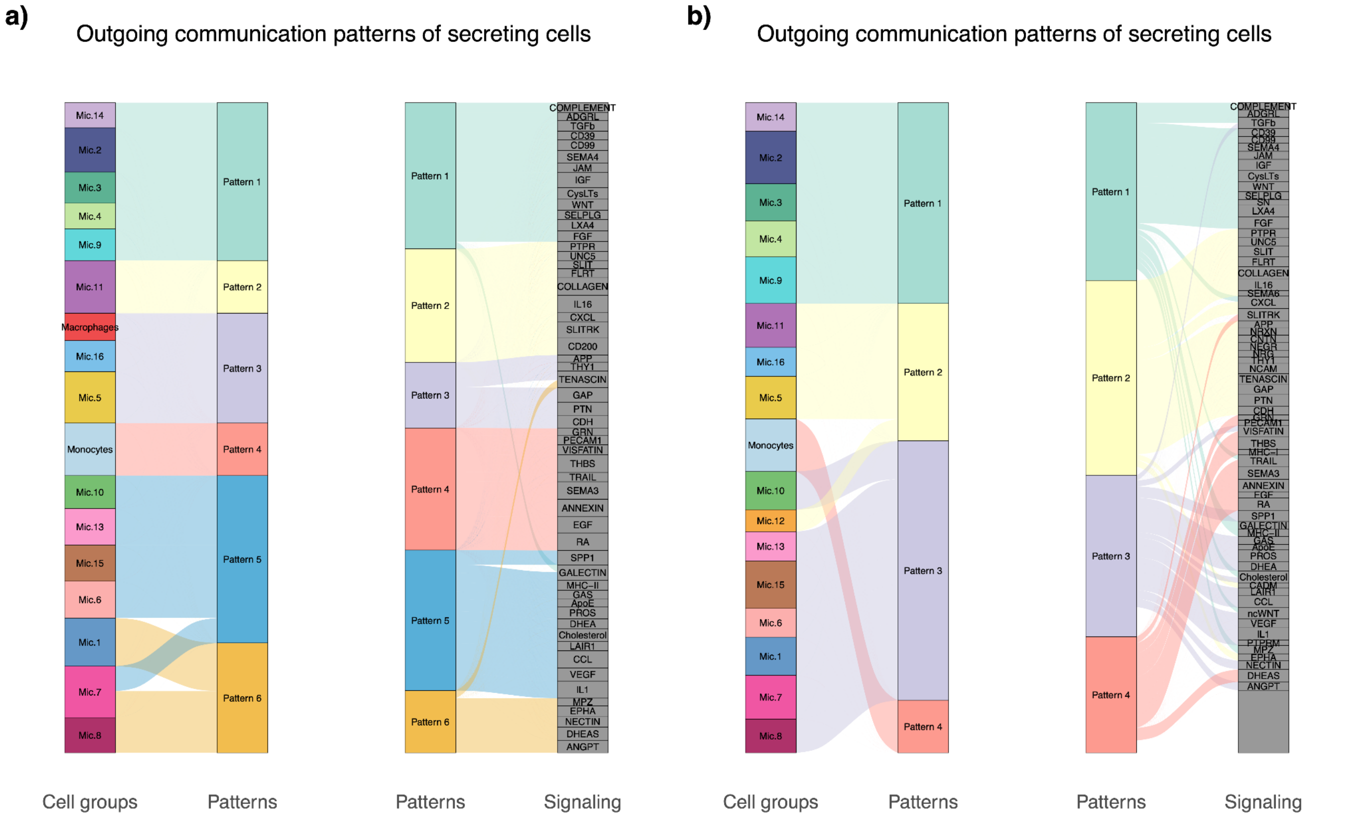


**Supplementary Figure 13: Outgoing patterns in CT vs AD.**

a) River plot of the most enriched cell groups and the signaling pathways associated with each inferred pattern in the exit signaling of the CT groups and, b) AD.

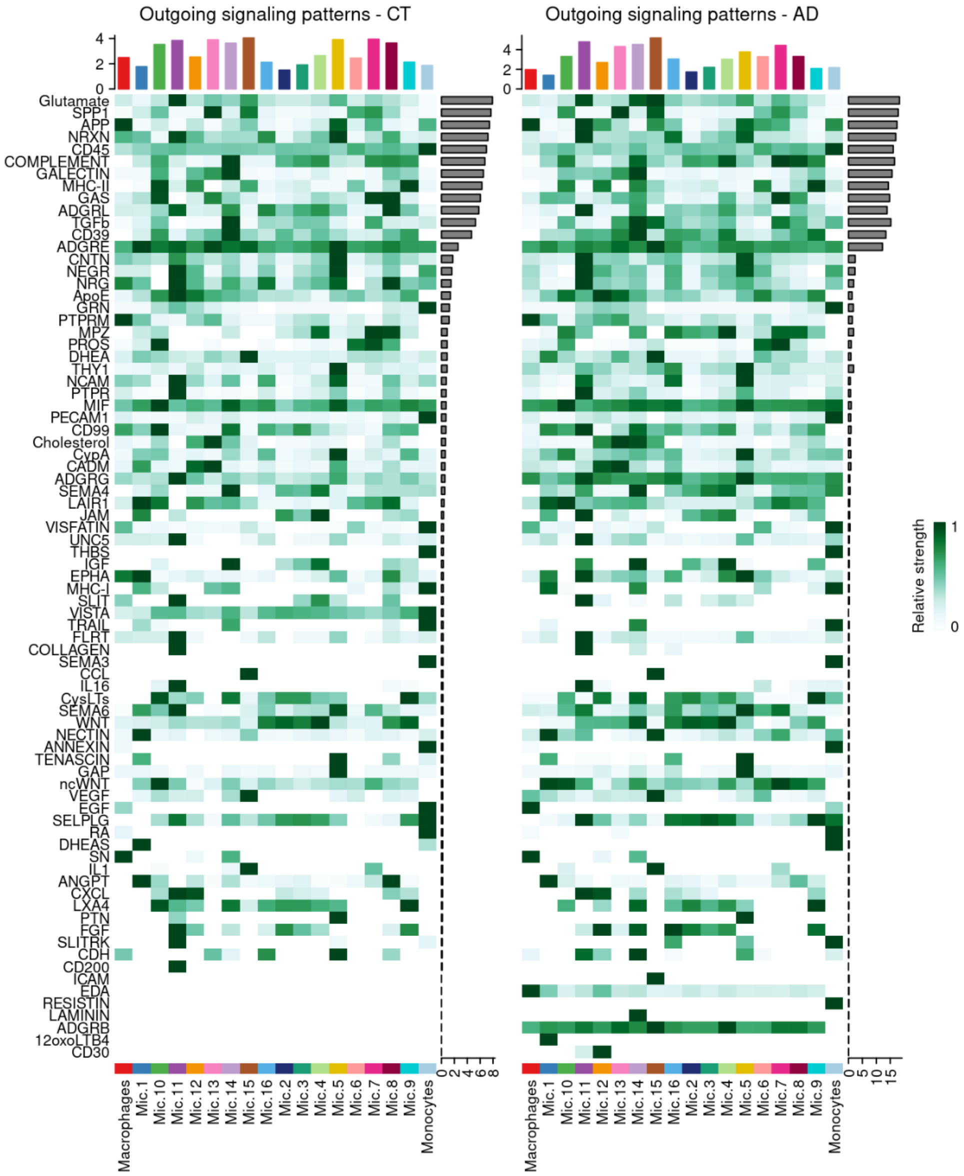


**Supplementary Figure 14: Outgoing signalling patterns CT vs AD.**
Heatmaps of the relative strength of signaling pathways in the outgoing signaling patterns in each cell group, in CT and, b) AD.

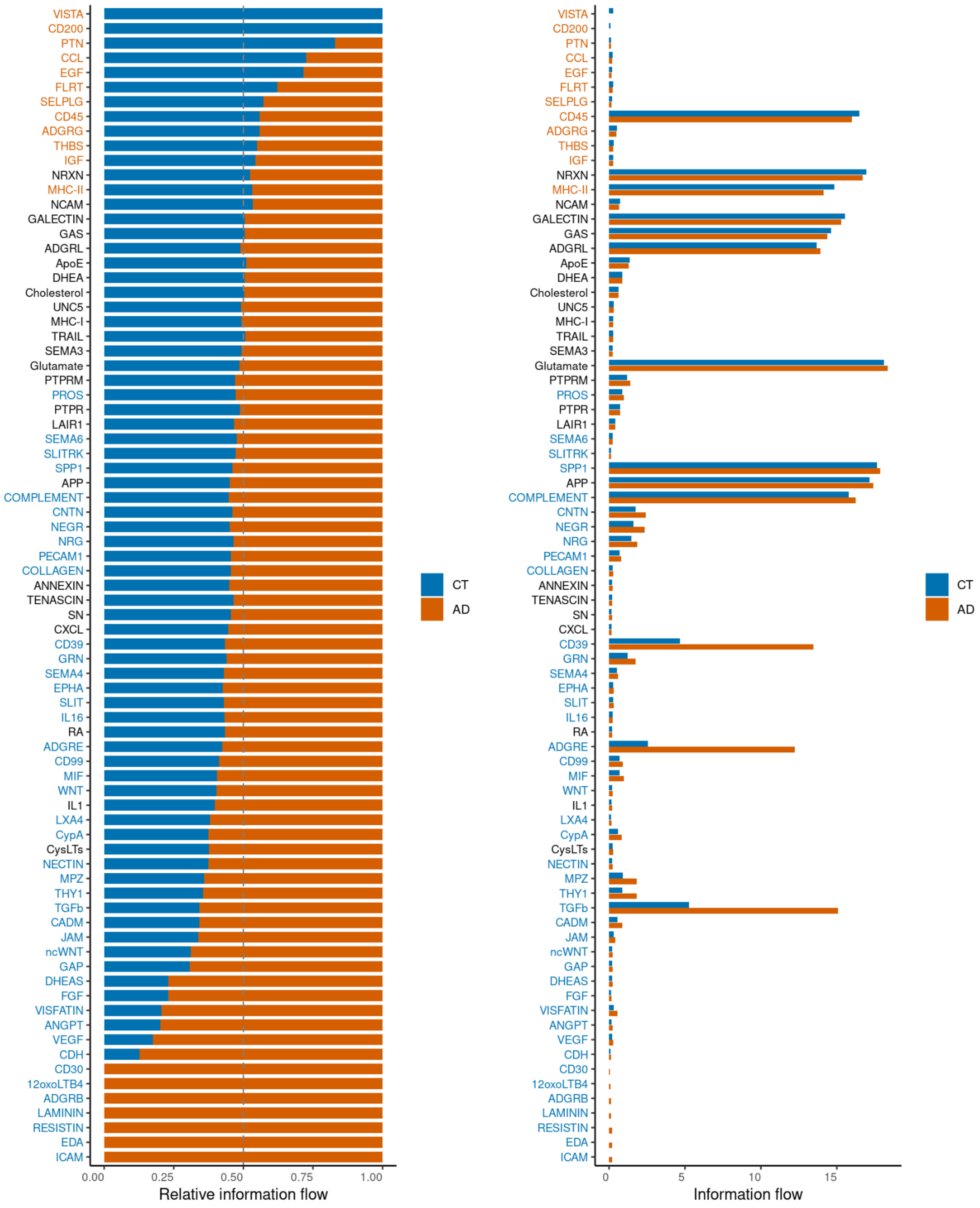


**Supplementary Figure 15: Relative information flow via CT and AD.**

Stacked bar chart of the signaling pathways flow in each group, applying the Wilcoxon test with a cutoff of 0.05. Colored labels indicate significant deviations in contribution (relative contribution - 1 > 0.05, P < 0.05): orange (enriched in AD), blue (enriched in CT), or black (no significant difference).

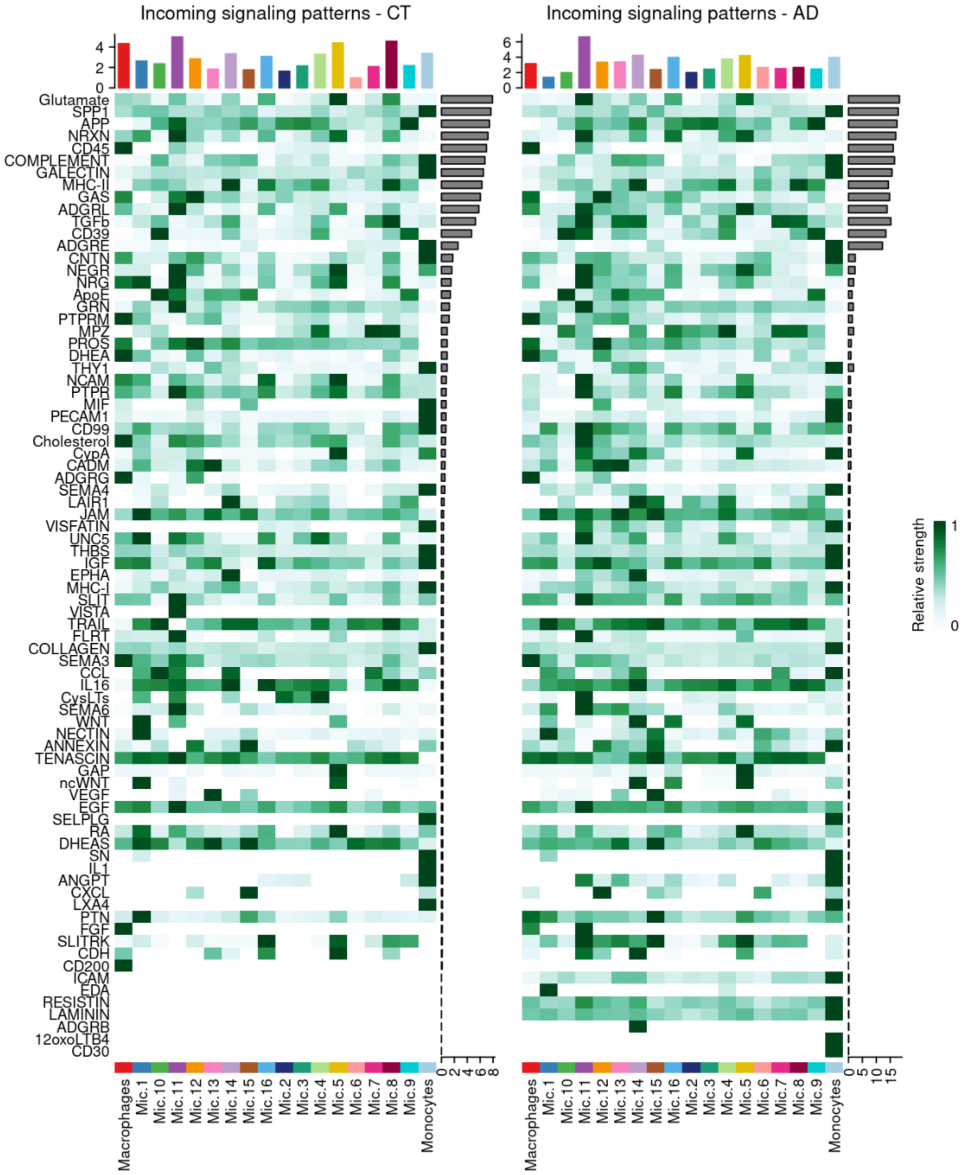


**Supplementary Figure 16: Incoming signaling patterns CT vs AD.**
Heatmaps of the relative strength of signaling pathways in the incoming signaling patterns in each cell group, in CT and, b) AD.

### Session 03: Replication analysis.

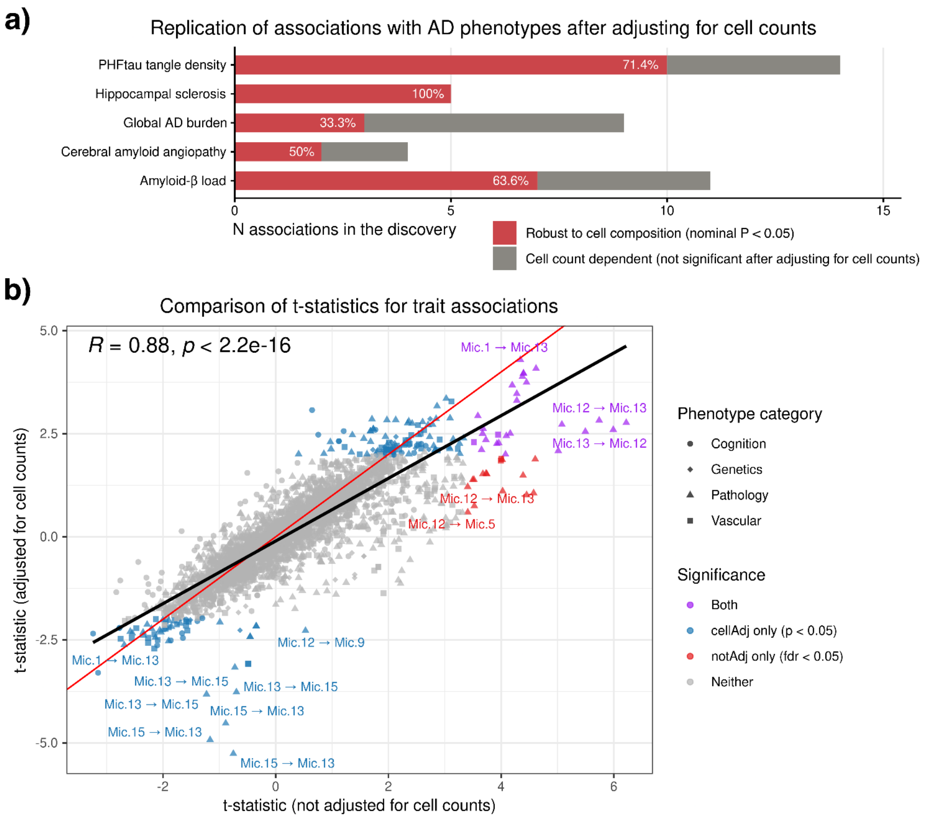


**Supplementary Figure 17: Replication of association analysis with AD phenotypes.**

a) Bar plot showing the percentage of associations related to Figure 4 that were replicated after adjusting for cell counts. b) Comparison of t-statistics for AD trait associations using the full summary statistics. The x-axis shows t-statistics results of regression models without adjustment for cell counts, and the y-axis shows t-statistics results after adjustment for cell counts. Each point is a pair of cell weights associated with a phenotype. Point shapes represent the phenotype category. Colors represent the significance relative to the model: purple = significant in both models; red = significant only in the not adjusted model; blue = significant only in the adjusted model; gray = not significant in either. For models not adjusted by cell counts, significance criteria was FDR corrected p-values (FDR < 0.05), while nominal significance (P < 0.05) was considered for models adjusted by cell counts. The Spearman correlation and respective p-value are shown on top. The regression line is shown in black, the diagonal is shown in red color.

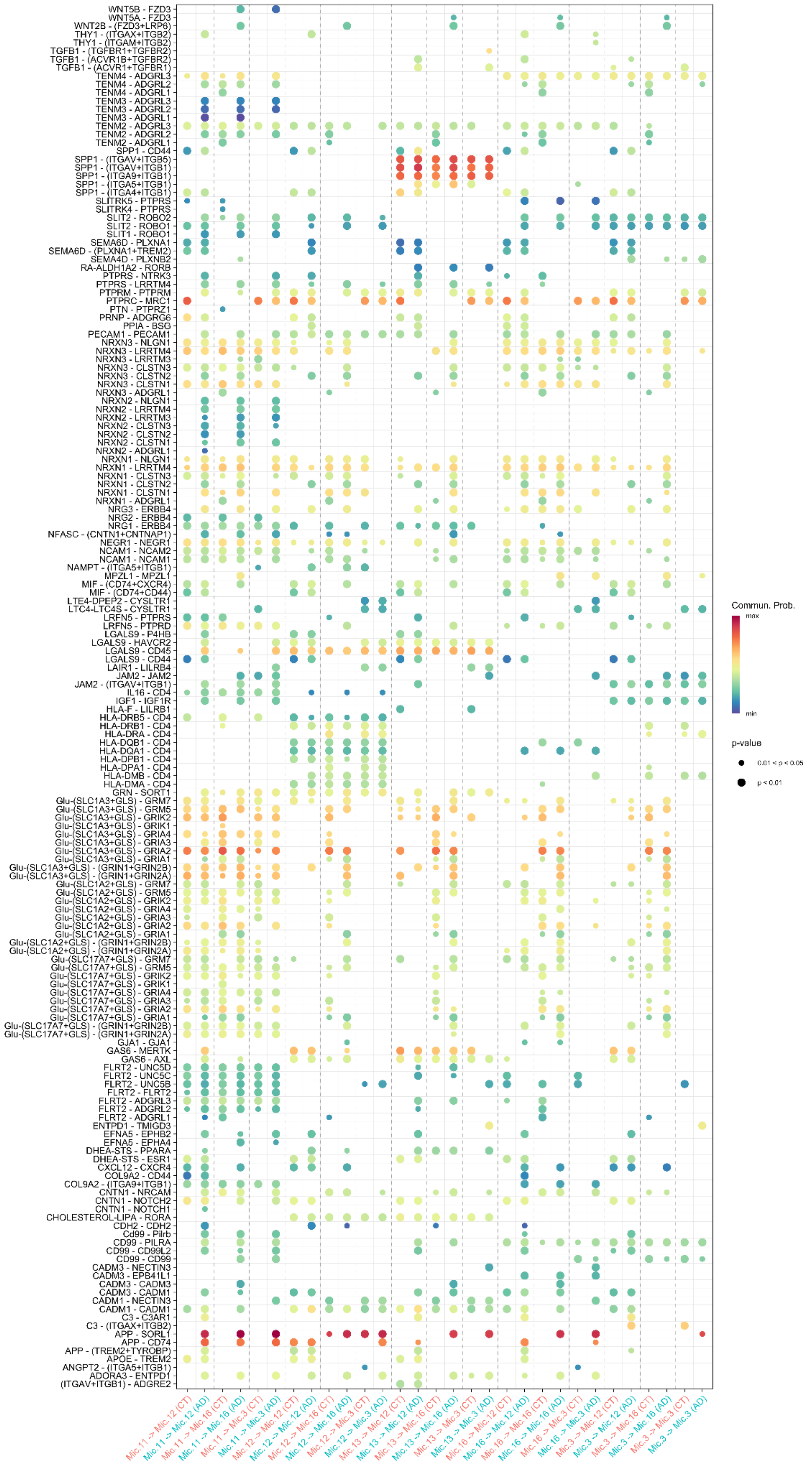


**Supplementary Figure 18: Dysfunctional signaling in CT vs AD.** Bubble plot of dysfunctional signaling. Interactions are compared between CT and AD through the probability of communication. Pairs are read as ligand-receptor; the + sign within the parentheses indicates a multimeric receptor complex. Each interaction shown occurs between the emitter groups Mic.11, Mic.12, Mic.13, Mic.16, and Mic.3 and the receiver groups Mic.12, Mic.16, and Mic.3.

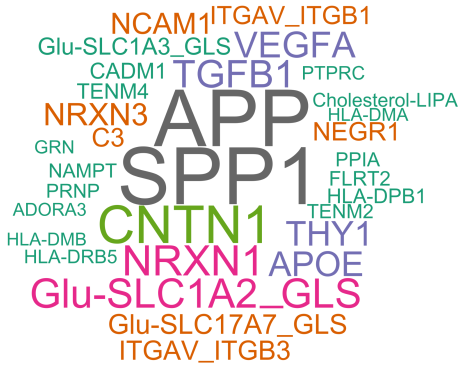


**Supplementary Figure 19: Word cloud.**
Ligands upregulated.

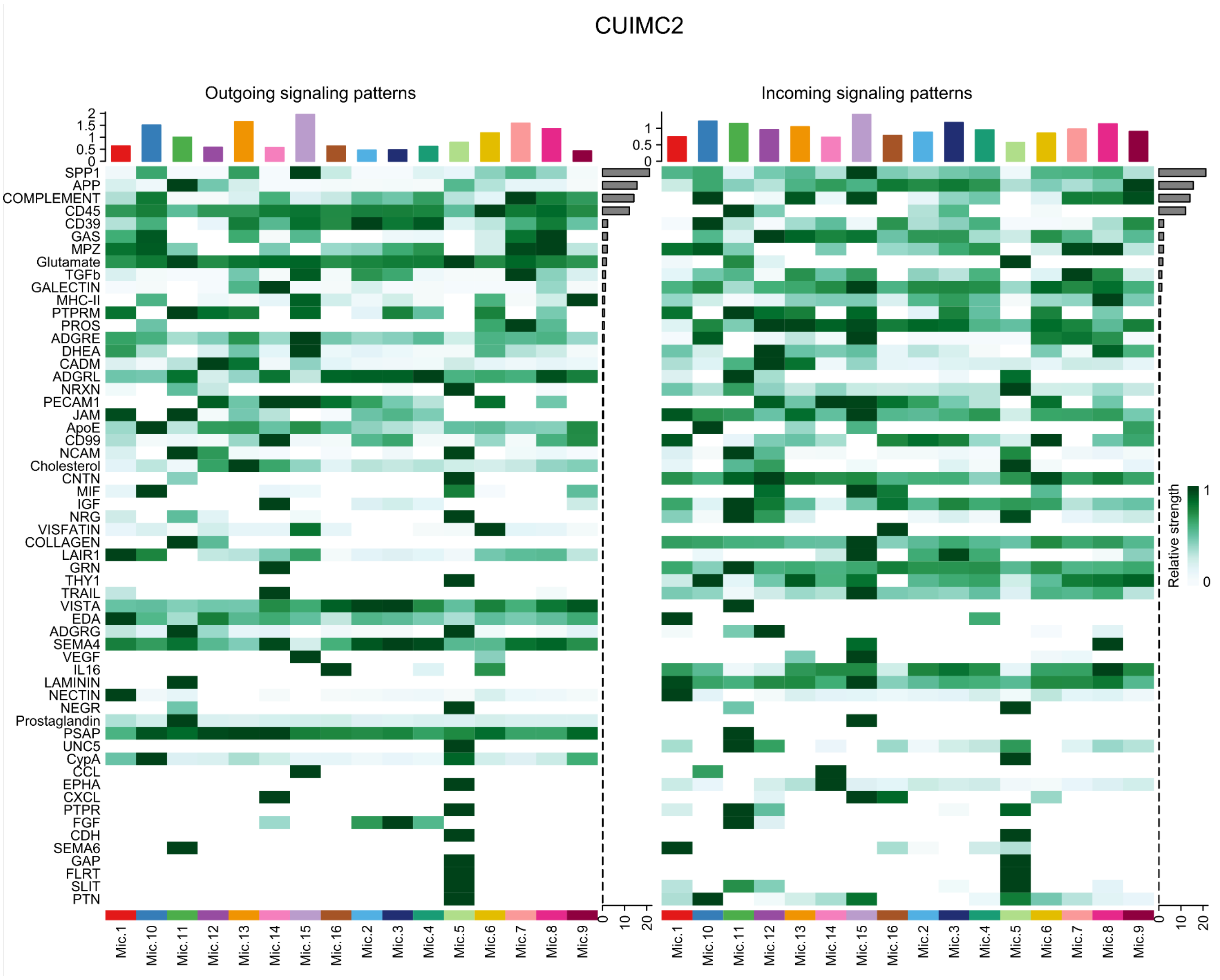


**Supplementary Figure 20: Signaling pathways of the CUIMC2.**

Heatmap of the relative strength of signaling pathways for each cell group in the CUIMC2 replication dataset.

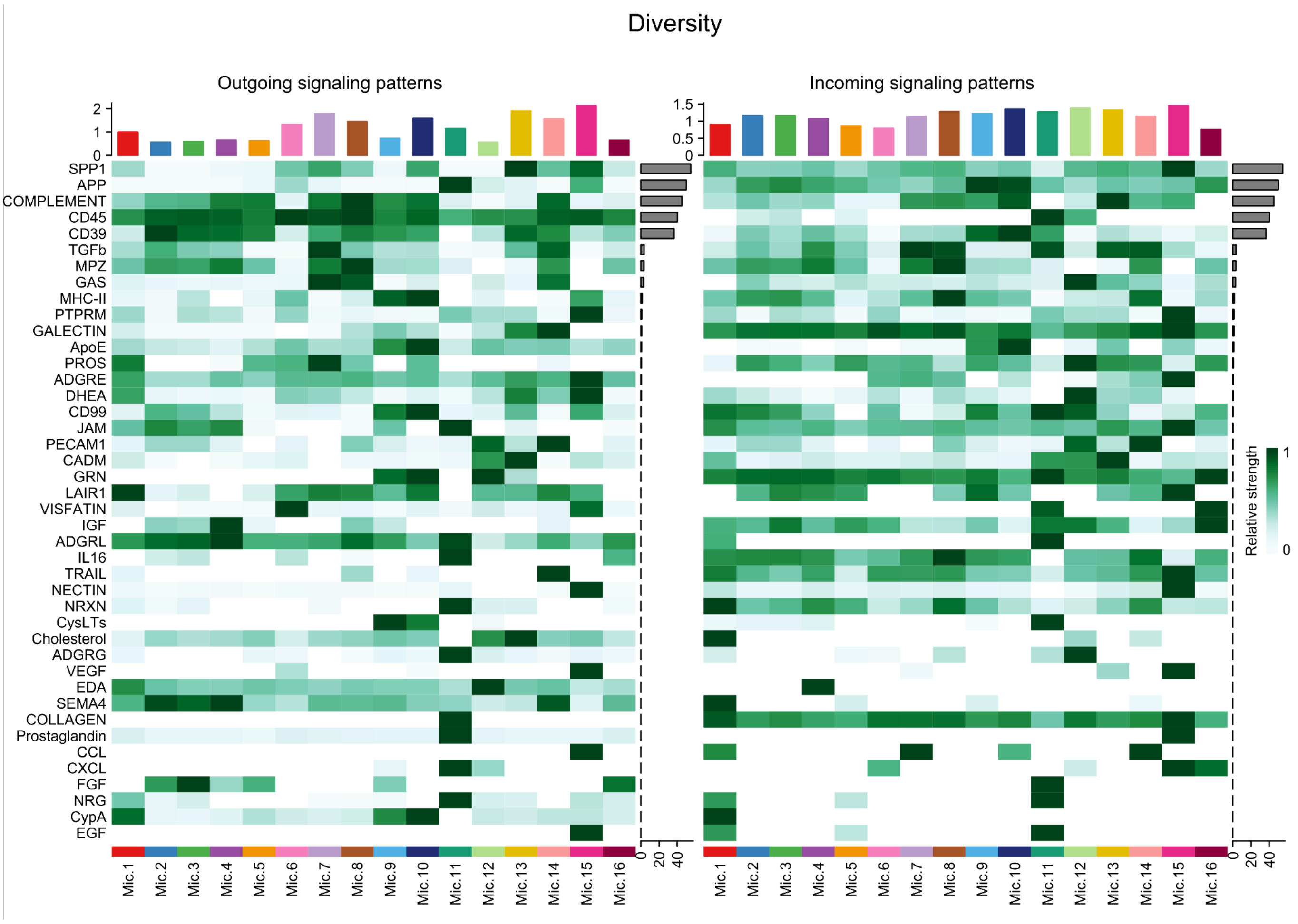


**Supplementary Figure 21: Signaling pathways of the Diversity dataset.**

Heatmap of the relative strength of signaling pathways for each cell group in the Diversity replication dataset.

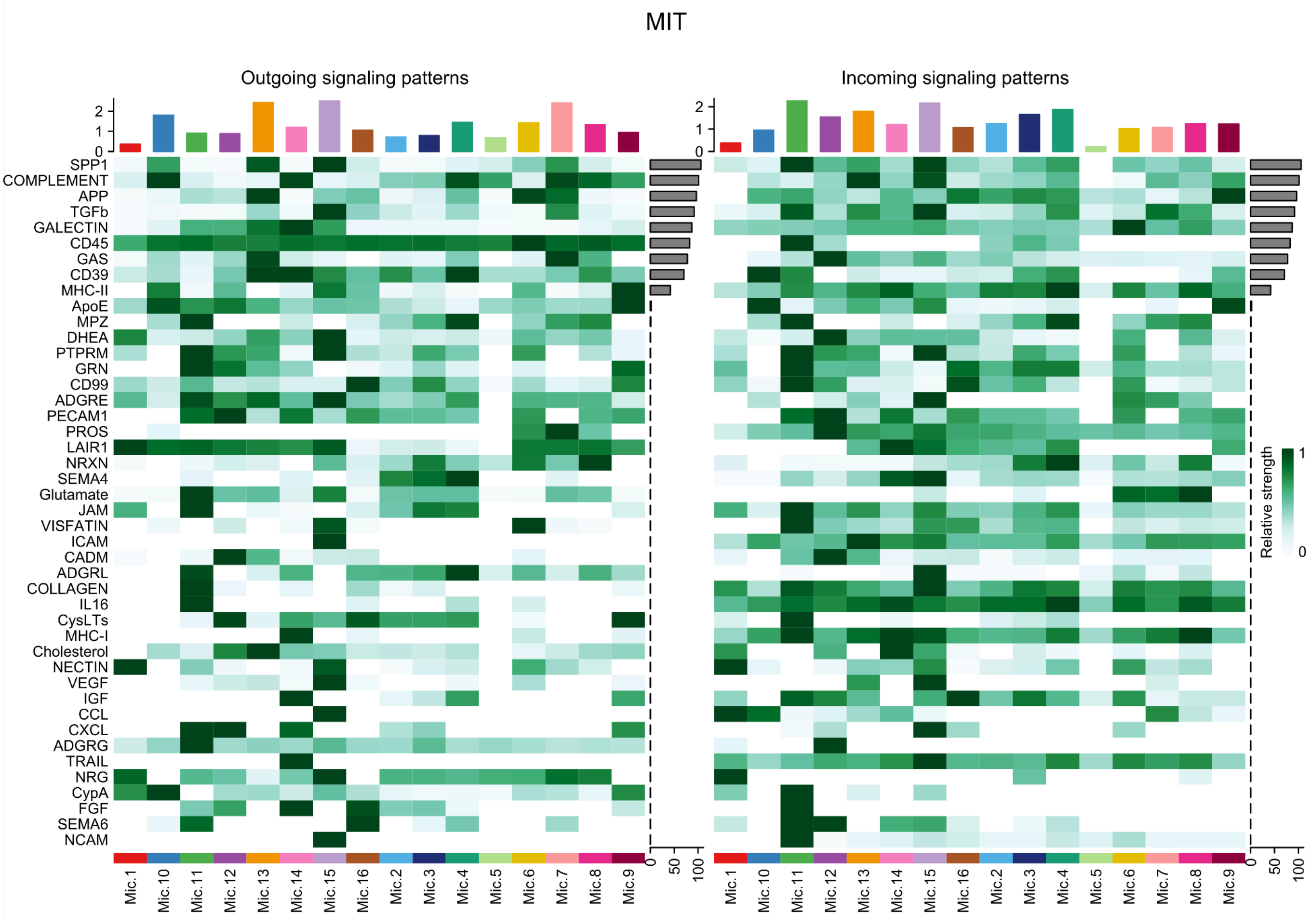


**Supplementary Figure 22: Signaling pathways of the MIT.**

Heatmap of the relative strength of signaling pathways for each cell group in the MIT replication dataset.

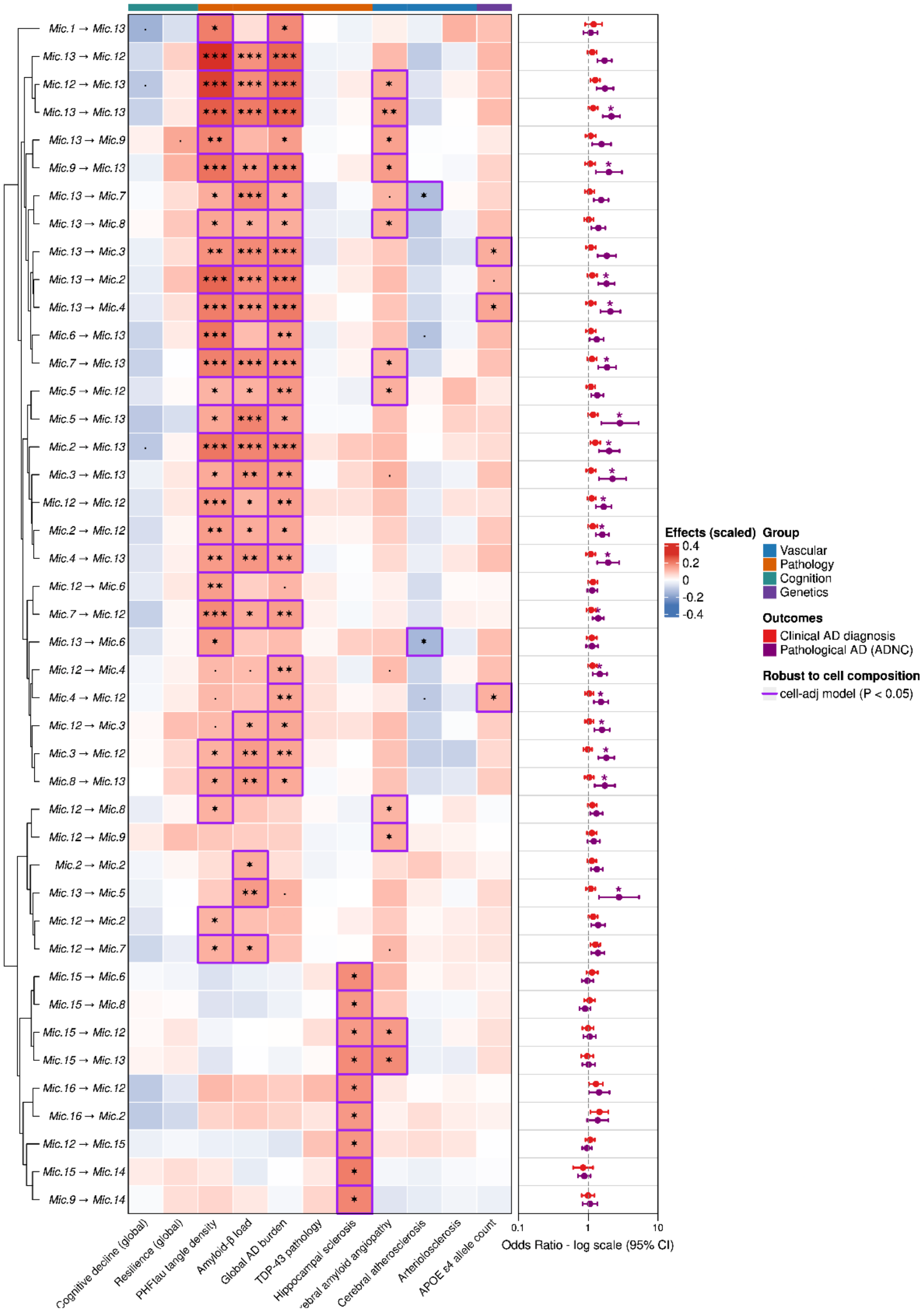


**Supplementary Figure 23: Association analysis with AD-related traits in datasets combined.**

Heatmap and forest plot of the associations from the Discovery dataset combined with the associations from the Replication datasets, totaling 840 participants. Associations of continuous and binary outcomes were measured via linear and logistic regression, respectively. In the heatmap, regression estimates are represented by color intensity and statistical significance showed as asterisks (FDR ***p<0.001, **p<0.01, *p<0.05). Phenotypes are labeled according to their category. Purple squares highlights results robust to cell composition adjustment. The forest plots show results for binary outcomes as log Odds Ratio (logOR) and 95% CI. Results nominally significant after cell composition adjustment are indicated with a purple asterisk.

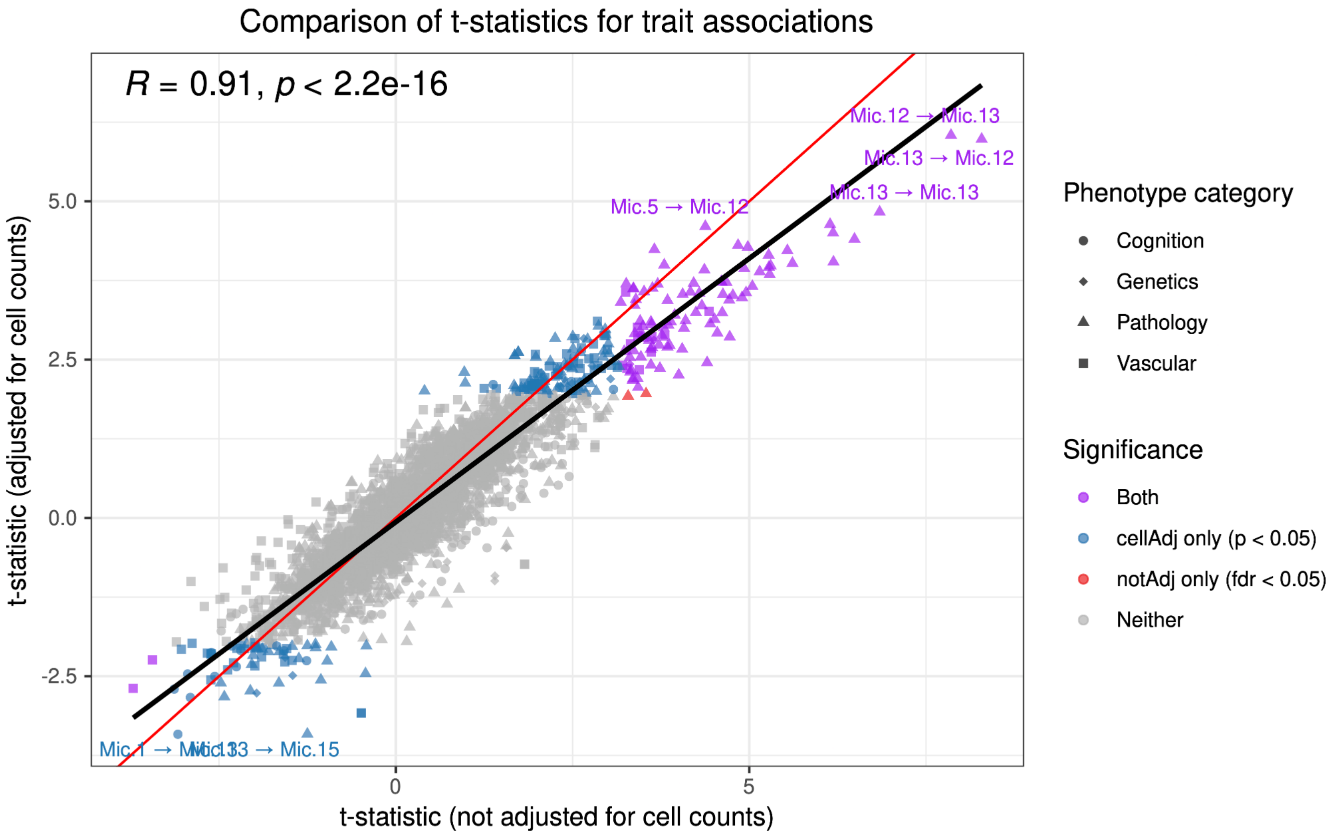


**Supplementary Figure 24: Comparison of AD-trait association models adjusted and not adjusted by cell counts using combined datasets.**

Data show results for the associations including the Discovery dataset combined the Replication datasets, totaling 840 participants. The x-axis shows t-statistics results of regression models without adjustment for cell counts, and the y-axis shows t-statistics results after adjustment for cell counts. Each point is a pair of cell weights associated with a phenotype. Point shapes represent the phenotype category. Colors represent the significance relative to the model: purple = significant in both models; red = significant only in the not adjusted model; blue = significant only in the adjusted model; gray = not significant in either. For models not adjusted by cell counts, significance criteria was FDR corrected p-values (FDR < 0.05), while nominal significance (P < 0.05) was considered for models adjusted by cell counts. The Spearman correlation and respective p-value are shown on top. The regression line is shown in black, the diagonal is shown in red color.
